# *CTHRC1*^+^ Fibroblasts and *SPP1*^+^ Macrophages Synergistically Contribute to Pro-Tumorigenic Tumor Microenvironment in Pancreatic Ductal Adenocarcinoma

**DOI:** 10.1101/2024.04.23.590663

**Authors:** Evan Li, Hoi Ching Cheung, Shuangge Ma

## Abstract

Pancreatic ductal adenocarcinoma (PDAC) is an extremely lethal cancer that accounts for over 90% of all pancreatic cancer cases. With a 5-year survival rate of only 13%, PDAC has proven to be extremely desmoplastic and immunosuppressive to most current therapies, including chemotherapy and surgical resection. In recent years, focus has shifted to understanding the tumor microenvironment (TME) around PDAC, enabling a greater understanding of biological pathways and intercellular interactions that can ultimately lead to potential for future drug targets. In this study, we leverage a combination of single-cell and spatial transcriptomics to further identify cellular populations and interactions within the highly heterogeneous TME. We demonstrate that *SPP1*^+^*APOE*^+^ tumor-associated macrophages (TAM) and *CTHRC1*^+^*GREM1*^+^ cancer-associated myofibroblasts (myCAF) not only act synergistically to promote an immune-suppressive TME through active extracellular matrix (ECM) deposition and epithelial mesenchymal transition (EMT), but are spatially colocalized and correlated, leading to worse prognosis. Our results highlight the crosstalk between stromal and myeloid cells as a significant area of study for future therapeutic targets to treat cancer.

## Introduction

About half a million new cases of pancreatic ductal adenocarcinoma are diagnosed globally every year, but only around 10% of patients overall survive. PDAC accounts for 450 thousand deaths globally, making it the 7th leading cause of cancer deaths [1]. In the US, the 5-year survival rate has recently exceeded 13% in 2024, making it one of the most lethal cancers [2]. Not only is PDAC extremely hard to detect before metastasis, but its heterogeneity among patients limits the effectiveness of current chemotherapies and immunotherapies. A key aspect of the complexity of PDAC lies in the tumor microenvironment, harboring a diverse population of cells including cancer-associated fibroblasts (CAF), tumor-associated macrophages, endothelial cells, neutrophils, and T and B lymphocytes. Despite the abundance of immune cells in close proximity to the tumor, the immunosuppressive nature of the TME greatly limits the immune response, leading to poor prognosis and difficulties in treatment.

The TME is also characterized by an extremely high percentage of extracellular matrix, consisting of collagens, integrins, and matrix metalloproteinases [3], acting as a physical barrier to immune infiltration. With an extensive population of fibroblasts that range from tumor-promoting and tumor-suppressing [4], two main populations of fibroblasts have been defined by Ohlund *et al.* in 2017: pro-fibrotic myofibroblasts (myCAF) characterized by α*-SMA* (smooth muscle actin), and pro-inflammatory fibroblasts (iCAF) expressing high levels of interleukins [5]. Although these two classifications reveal distinct roles of CAF in matrix remodeling, growth factor secretion, and tumor-stimulation [6], the heterogeneity of PDAC has revealed the need to identify more specific groups and interactions. Since then, many additional subtypes of CAF have been identified using single-cell transcriptomics, and a study by Cords et al. [7] identified 7 more additional subtypes of CAF in breast cancer with various functions that included angiogenesis, antigen-presentation, and interferon production. The roles of CAF in tumors are therefore complex and cannot be understated. However, as the targeting of myCAF and knockout of α-*SMA* in PDAC has led to tumor indifferentiation, leading to a worse prognosis in both humans and mice [8], a multi-faceted approach is needed in order to target the stroma. In recent years, attention has been increasing on the crosstalk between pro-fibrosis TAM and CAF in particular.

Macrophages play a pivotal role in orchestrating immunosuppression in the TME through diverse mechanisms. They can be generally classified into M1 anti-tumor and M2 pro-tumor [9], but this system does not show the heterogenic role of TAM in detail. It is widely known that anti-inflammatory and pro-fibrotic M2 TAM secrete various immunosuppressive cytokines and chemokines, along with transforming growth factor-beta (*TGF-β*) to promote fibrosis, EMT, and foster the recruitment and activation of CAF [10,11]. M2 TAM have also been shown to promote cell proliferation, angiogenesis, and phagocytosis to reduce inflammation [9]. High expression of these TAM are also correlated to high expression of *PD-L1* and resistance to anti-*PD-L1* treatment, a therapy that has shown to not be effective in PDAC in part due to its non-immunogenic nature [12,13]. Recently, *SPP1* (osteopontin) has been associated as a hallmark of pro-tumor macrophages [14], and their functions in the TME have been well established. It is noteworthy that *SPP1^+^* macrophages have been implicated in activation of myofibroblasts in other immune mediated diseases, such as kidney and lung fibrosis [15]. However, the interactions between *SPP1*^+^ TAM and other cells in the TME has not been fully investigated.

In this study, we unveil a distinct population of TAM characterized by *SPP1* and *APOE* along with another population of myCAF expressing *CTHRC1* and *GREM1*. Using single cell transcriptomics, we establish a high degree of correlation, highlighting their contributions in immune-suppressive capacities, driving ECM deposition, facilitating matrix remodeling, and EMT, consequently fueling tumor progression. We validate results with a spatial approach, demonstrating the proximity and concordance of these two populations, reinforcing the significance of our results. This research serves as additional insight into the complex crosstalk between macrophage and stromal populations within the PDAC TME.

## Results

### scRNAseq Data Integration Reveals Diverse Tumor Microenvironment

To conduct this study, tumor samples (n=51), metastatic samples (n=6), and adjacent normal tissue (n=6) from a small subset of patients were obtained from four public scRNAseq datasets. After filtering for low quality cells or genes, a total population of 98,749 cells and 35,677 genes remained for downstream analysis, revealing an atlas of the TME (Fig. 1A). The major clusters were annotated according to canonical marker genes (Fig. 1B-C, Supplementary Fig. 1A): T cells (n = 27,862) were positive for *CD3E*, *CD4*, *CD8A*, and *GZMB*; B-cells (n = 5,071) expressed *MS4A1* and *CD79A*; plasma B (n = 1,516) had high expression of *CD79A* and *MZB1*; acinar cells (n= 1,694) exhibited *PRSS1* and *REG1A*, epithelial cells (n=29,781) expressed *EPCAM* and *KRT8*; endothelial (n = 1,803) showed *PECAM1* and *CDH5*; mast cells (n = 2,010) were positive for *TPSAB*1, myeloid cells (n= 21,645) expressed *CD16b*, *CD68*, and *CD14*; and stromal cells (n= 5,564) were defined by *COL1A1* and *LUM*. We then combined epithelial with acinar cells, mast cells with myeloid cells, B cells with plasma B cells, and endothelial cells with stromal cells to form five major groups for downstream analysis, with calculated total counts (Fig. 1D) and patient type composition (Fig. 1E), revealing a diverse tumor population composed primarily of epithelial, T-cells, and myeloid cells. These preliminary results show the abundance of immune cells in the TME; however, the poor prognosis and failures of current immunotherapies suggest mechanisms that prevent an effective immune response.

**Figure 1:**
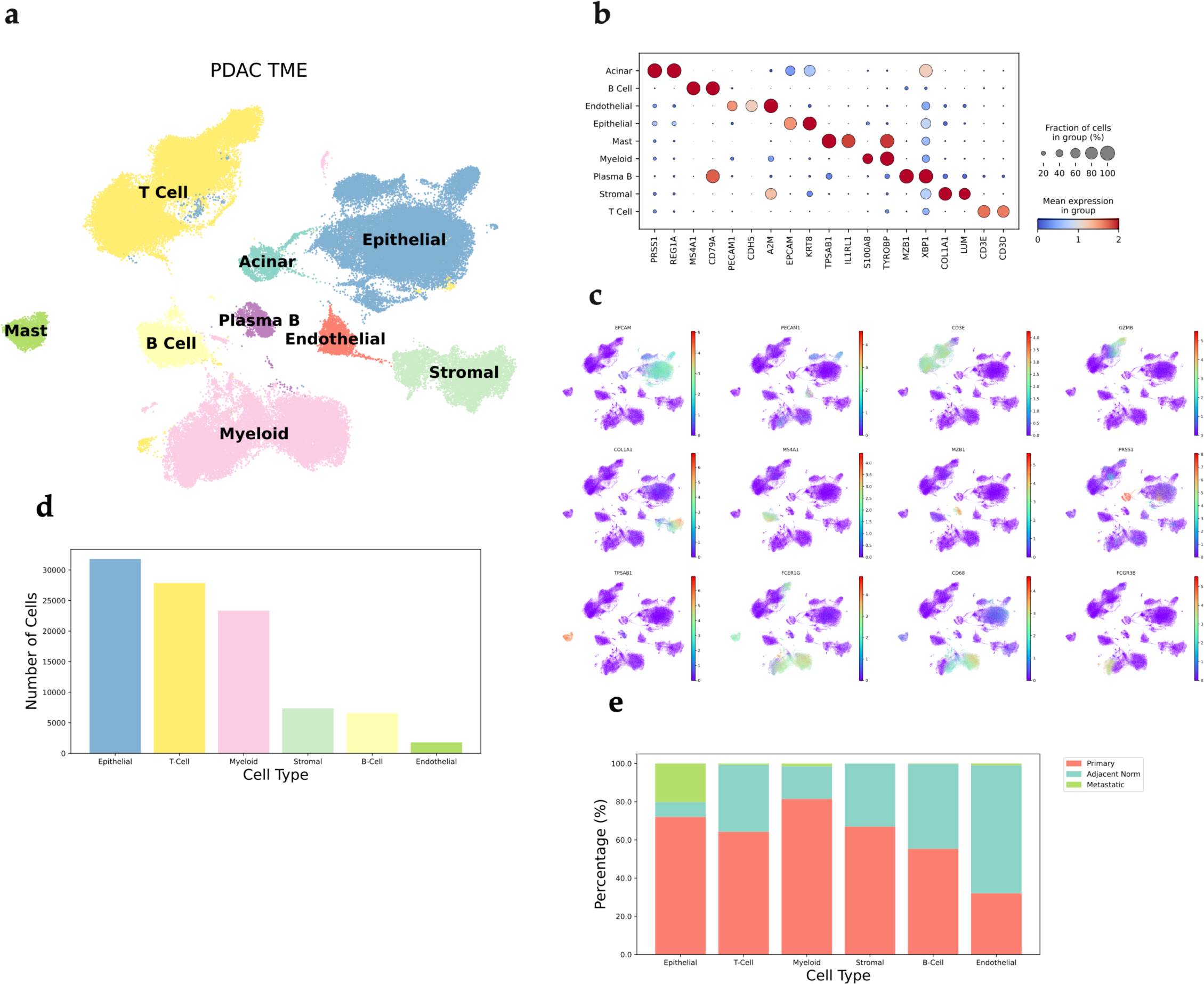
Single cell RNA-Seq reveals PDAC TME landscape. Canonical Cell Markers in PDAC TME. (A) Pipeline of scRNAseq analysis. (B) Uniform Manifold Approximation Projection (UMAP) of the Comprehensive Tumor Microenvironment, with a total of 63 samples, 98,749 cells and 35,677 genes. (C) Dot plot of canonical cell marker genes used to annotate major cells. (D) Bar plot of number of cells per cell type. (E) Bar plot of percentages of each condition per cell type.

### Categorization of T-Cells in the PDAC TME

First, we investigated the composition of T cells in the TME, as they are key parts of the adaptive immunity response against tumors. We labeled 11 groups under the general classification of CD4*^+^* / CD8*^+^* T-cells and *GZMB^+^* NK cells, and used reported genes in literature to identify specific T cell subtypes (Fig. 2A-B). CD4*^+^* T cells were comprehensively found in greater proportions in tumor populations (Fig. 2C), with CD4*^+^*Tcm (Diff = 7.2%, p = 0.031), FOXP3^+^ Treg (Diff = 7.6%, p = 0.035), Naive CD4*^+^* (Diff = 13.2%, p = 0.0021), and Th2 (Diff = 3.6%, p = 0.0059) all enriched. Populations of FOXP3^+^ Treg have been confirmed mediators in angiogenesis and immune-suppressive functions, contributing to the pro-tumor TME [16], although a recent study stated the role of Th2 cells in anti-tumorigenic responses [17].

**Figure 2:**
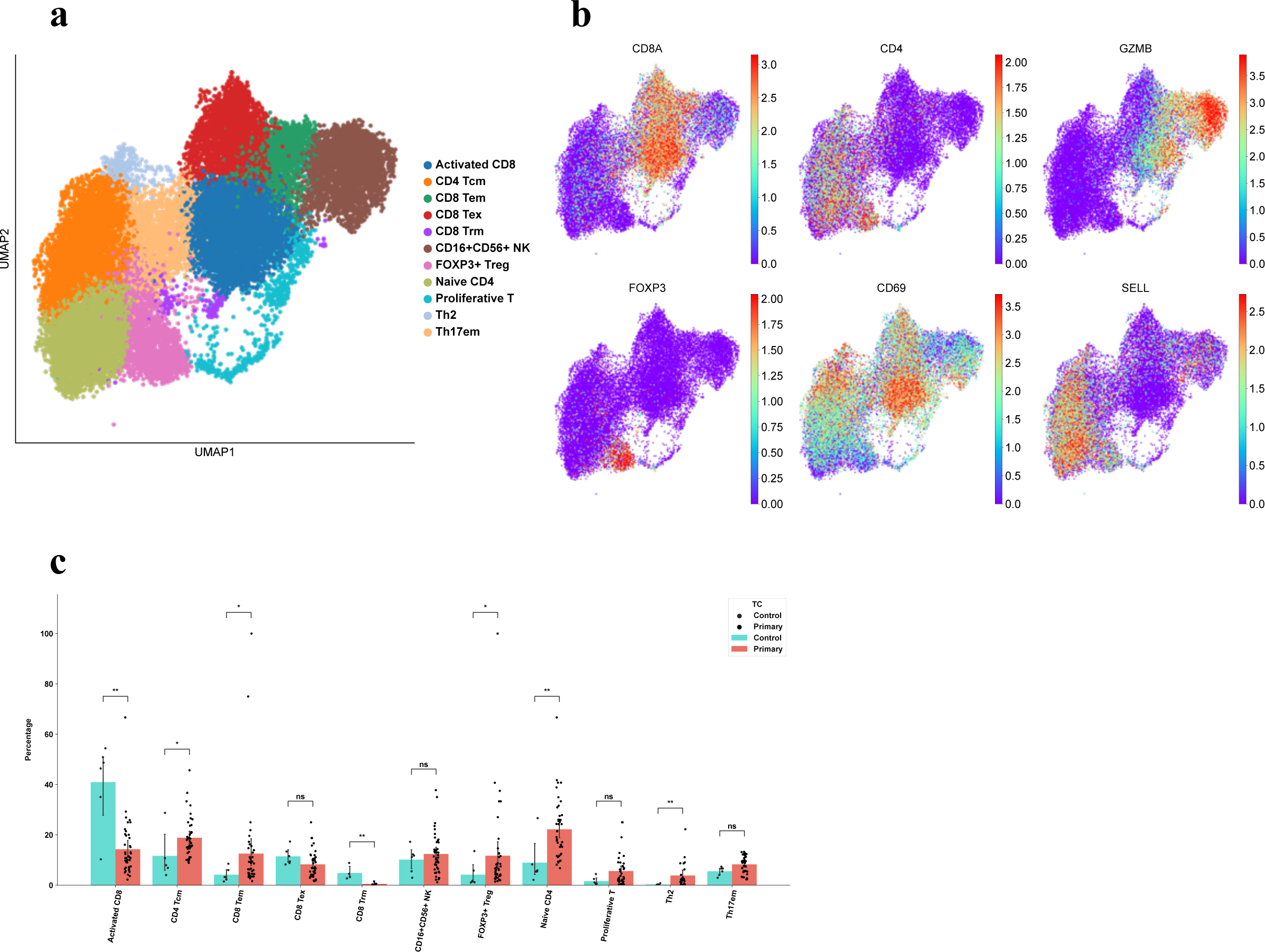
Characterization of T-Cells in the TME. Proportions of T cells in TME Underscore Low CD8^+^ T cell Infiltration and Activation. (A) UMAP characterization of T cell populations in the TME. (B) UMAP of defining marker genes in T-cells populations. (C) Bar plot of percentage per group per patient. *CD4*^+^ Tcm (p = 0.031), FOXP3^+^ Treg (p = 0.035), Naive CD4^+^ (p = 0.0021), Th2 (p = 0.0059), activated CD8^+^ (p =0.0026), and CD8^+^ Trm (p = 0.0095) are statistically significant using Mann-Whitney U Test.

Conversely, the majority of CD8^+^ T cells were found in lower levels in tumor samples, with activated CD8*^+^* (Diff = 26.7%, p =0.0026) and CD8^+^ Trm (Diff = 4.3%, p = 0.0095) being significant in adjacent normal tissue (Fig. 2C). Although studies have logically shown the increased presence of exhausted CD8*^+^* T cells in tumors due to the immunosuppressive environment [18,19], we found no significant difference in T exhausted proportions between adjacent and tumor (Diff = 3.2%, p = 0.088), with even activated CD8*^+^* cells in the tumor surpassing the T exhausted proportion (Fig. 2C). This suggests that the infiltration and effectiveness of CD8*^+^* T cells in the tumor is extremely low, hence its classification as a “cold tumor” owing to the failures and challenges of anti-*PDL1* therapies in PDAC [20]. The increased number of CD4*^+^* regulatory and reduced CD8^+^ anti-tumor responses indicate the prevalent pro-tumor mechanisms in the TME, and more research is needed to uncover these biological processes.

### Characterization of Stromal Cells in the TME

As stromal cells have been known to contribute to an immunosuppressive TME while playing a critical role in the production of extracellular matrix in PDAC [21], we decided to investigate this cell type further to classify specific identities of these populations. Altogether, 12 populations of stromal cells were identified based on previously described gene markers (Fig 3A, Supplementary Fig. 2A), including *PECAM1^+^* endothelial cells (n = 1803). Smooth muscle cells (n = 1274) and pericytes (n = 523) were classified by high expression of *MYL9* and *α-SMA* (*ACTA2*), which suggests that *α-SMA* may not be suitable as a hallmark for myCAF, given their high expression across multiple groups. We therefore identified our large subset of myCAF based on their expression of matrix-associated genes (collagens, proteoglycans, and matrix metalloproteinases) and *CTHRC1*, which were then further divided into two groups: canonical myCAF (n = 444) with high *CTHRC1* expression, and *CTHRC1^+^GREM1^+^*myCAF (n = 1413) which were also positive for *α-SMA* (Supplementary Fig. 2B). *CLU^+^* fibroblasts (n = 515) resembled a smooth muscle phenotype with expression of *CLU* and *ADIRF*. A large population of fibroblasts expressed the complements *C3* and *C7*, suggesting their proinflammatory nature; we differentiated them into three subtypes, *C3^+^RARRES1^+^* CAF (n = 873), *C3^+^SFRP1^+^* CAF (n = 264), and a group that exhibited both an inflammatory and myofibroblast signature, which we designated *C3^+^CTHRC1^+^* CAF (n = 1413). Mesothelial cells acquired high expression of *KRT8* and *KRT18* (n = 90). A small subset of fibroblasts were shown to be antigen-presenting fibroblasts based on expression of *MHC-II* and *RGS5* (n = 78), while another group expressed *MKI67*, represented as proliferative CAF (n = 30).

**Figure 3:**
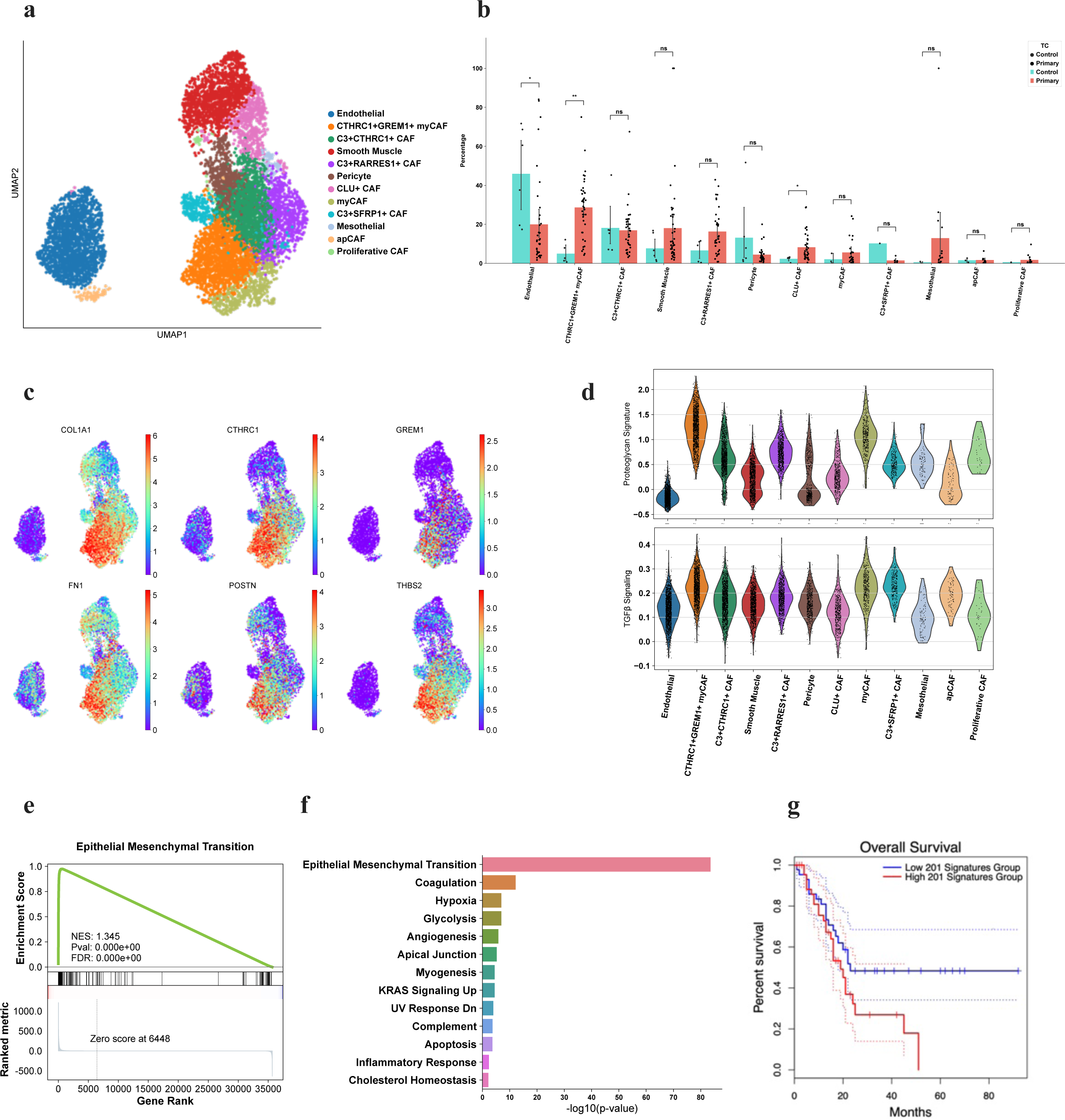
Characterization of Stromal cells reveals enriched tumor population of CTHRC1+GREM1+ myofibroblasts. *CTHRC1^+^GREM1^+^* myCAF contribute to fibrosis and pro-tumorigenic ECM production and remodeling. (A) UMAP characterization of stromal cell populations in the TME. (B) Bar-plot of percentage per group per patient. *CTHRC1^+^GREM1^+^* myCAF (p = 0.0014) and *CLU^+^*CAF (p = 0.025) are statistically significant using Mann-Whitney U-Test. (C) UMAP of DE genes in *CTHRC1^+^GREM1*^+^ myCAF. (D) Violin plots of proteoglycan and TGF-β signaling enrichment signatures. (E) GSEA plot of EMT pathway in *CTHRC1^+^GREM1^+^* myCAF. (F) Bar plot of upregulated pathways in *CTHRC1^+^GREM1^+^* myCAF using Enrichr. (G) Kaplan-Meier survival curve analysis of *CTHRC1^+^GREM1^+^*myCAF (TCGA cohort).

### *CTHRC1^+^GREM1^+^* myCAF Contribute to Fibrosis, Epithelial Mesenchymal Transition and is Linked to Poor Survival

Two groups, *CTHRC1^+^GREM1^+^* myCAF (Diff = 23.7%, p= 0.0014) and *CLU^+^* CAF (Diff = 5.9%, p = 0.025) were found in significantly higher proportions in tumor patients (Fig. 3B), while other groups such as smooth muscle (Diff = 10.4%), *C3^+^RARRES1^+^*CAF (Diff = 9.7%), and canonical myCAF (Diff = 3.5%) were trending towards tumor-enriched. In particular, *CTHRC1^+^GREM1^+^* myCAF expressed high levels of collagens (Fig. 3C, Supplementary Fig. 2B-C), including type-1 (*COL1A1*, *COL1A6*), type-3 (*COL3A1*), type-5 (*COL5A2*), and type-6 (*COL6A1*), which contribute to tumor cell migration and proliferation through collagen/integrin interactions [22,23]. These fibroblasts also expressed matrix metalloproteinases *MMP2*, *MMP11* and *MMP14* (Supplementary Fig 2B), which are not only key components of matrix remodeling [24], but lead to cancer progression and worse survival in patients [25,26]. We also observed increased expression of proteoglycans such as *CTHRC1* and *THBS2* that were unique to myCAF (Fig. 3C-D), which have both been determined to contribute to EMT through the *Wnt/b-catenin* and *NF-kB* pathway, leading to tumor invasiveness and poor survival [27,28,29]. In line with these findings, expression of *CTHRC1* was upregulated in tumors from all cancers in the TCGA cohort (Supplementary Fig 2D), suggesting these ECM protein networks are not limited to PDAC. *FN1* was also an important gene in *CTHRC1^+^GREM1^+^* myCAF (Fig. 3C), which has been shown to promote angiogenesis and metastasis of tumor cells through integrin signaling, leading to the activation of the *FAK* pathway and also contributing to EMT [30]. Interestingly, *ITGB1* and *ITGB5* were discovered to be present on these myCAF (Supplementary Fig. 2B), implying the role of these two integrins as a vital form of crosstalk between myCAF and ECM proteins (Supplementary Fig. 2E) [31,32].

To elucidate the biological pathways of this fibroblast phenotype, gene set enrichment analysis revealed that *CTHRC1^+^GREM1^+^*myCAF contributed towards increased levels of *TGF-β* signaling (Fig. 3D). Further analysis revealed that EMT was an extremely significant mechanism regulated by *CTHRC1^+^GREM1^+^* myCAF (Fig. 3E-F, Supplementary Fig. 2F), suggesting these fibroblasts support cancerous cell differentiation and proliferation leading to rapid tumor growth and potential metastasis. In addition, these cells were also found to help promote hypoxia (*ANXA2, SDC2, LOX*) and angiogenesis (*POSTN, VCAN, LUM*) (Fig. 3F). To evaluate the clinical effect on patients, we extended our study to include the TCGA cohort, finding that patients with a high signature of *CTHRC1^+^GREM1^+^*myCAF were shown to have significantly worse survival, implicating these cells as not only pro-fibrotic but pro-tumorigenic (Fig. 3G).

### Characterization of Myeloid Cells in the TME

Although fibroblasts are the main contributor to ECM deposition and remodeling, the function of myeloid cells as crucial components of the pro-tumorigenic TME in PDAC reveals the need to investigate these complex interactions further [33]. Altogether, we classified our myeloid subset into 15 groups (Fig. 4A, Supplementary Fig. 3A), including a known population of *TPSAB1^+^* mast cells (n = 2010). Neutrophils were categorized into three groups, with high expression of *CD16b* and *CD62L* throughout all cells; the first expressed high levels of interferons (*IFIT2* and *IFIT3*) which were labeled as *IFN^+^*neutrophils (n = 1647). Another group expressed high levels of *MMP9* and *MMP25*, which we labeled as *MMP9^+^* neutrophils (n = 356), while the last group was called *GMFG^+^* neutrophils (n = 2610). Four groups of monocytes were also found, which were labeled as *CD14^+^CD16^-^*monocytes (n = 1337), *CD16*^+^ monocytes (n = 440), *ITGB2*^+^ monocytes (n = 471), and *IL1B^+^* monocytes (n = 2880). A main group of classical dendritic cells (cDC) expressed high levels of *MHC-II* markers and *CD74* (n = 1856), and interestingly a small population of plasmacytoid DC were also noted by their expression of *CLEC4C* and *IL3RA* (n = 148). Macrophage populations characterized by *CD68* expression composed of the largest proportion of total myeloid cells, which were labeled into four groups: monocyte-like macrophages expressed *CD16a* and *MHC-II* (n = 2256), *C1Q*-high macrophages expressed *C1QA* and *C1QB* (n = 2748), *SPP1^+^APOE^+^*macrophages (n = 3226), and *SPP1^+^VEGFA^+^* macrophages (n = 1320). A small group of proliferating macrophages expressing *MKI67* (n = 176) were also identified.

**Figure 4:**
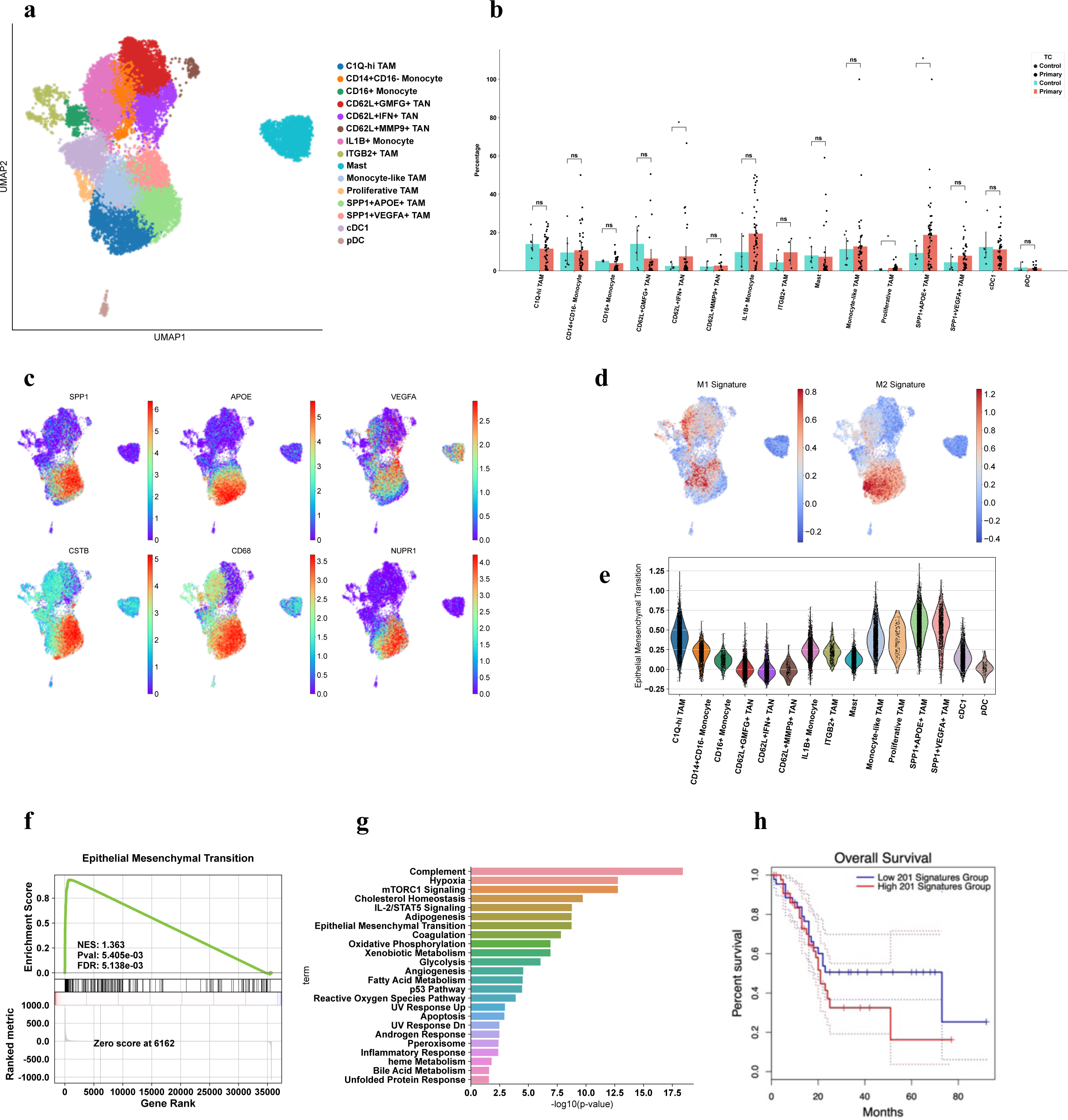
Characterization of myeloid cells in TME reveals enriched population of SPP1+ macrophages. *SPP1^+^APOE^+^* TAM exhibit pro-fibrosis and pro-tumor properties. (A) UMAP characterization of myeloid cell populations in the TME. (B) Bar plot of percentage per group per patient. *SPP1^+^APOE^+^*TAM (p = 0.011) and *IFN^+^* (p = 0.046) TAN are statistically significant using Welch’s T-Test. (C) UMAP plots of significant DE genes in *SPP1^+^APOE^+^* TAM. (D) UMAP plots of M1 and M2 macrophage signatures. (E) Violin plot of EMT enrichment signature. (F) GSEA plot of EMT pathway. (G) Bar plot of upregulated pathways in *SPP1^+^APOE^+^*TAM using Enrichr. (H) Kaplan-Meier survival curve for *SPP1^+^APOE^+^*TAM (TCGA cohort).

### *SPP1^+^* TAMs Enriched in Tumor and Contribute to Pro-Tumorigenic Functions and Lead to Worse Survival

We compared proportions of each myeloid subtype in each patient, finding that *SPP1^+^APOE^+^* TAM (Diff = 9.5%, p = 0.011) and *CD62L^+^IFN*-high neutrophils (Diff = 5.0%, p = 0.046) were significantly enriched in tumor patients versus control (Fig. 4B). *SPP1^+^VEGFA^+^* TAM were also trending towards increased proportions in tumor samples (Diff = 3.4%, p-value = 0.21). All *SPP1^+^* TAM exhibited the M2-marker *CD68* with other marker genes such as CSTB and NUPR1 (Fig. 4C), as well as being enriched for M2 signature under the M1 and M2 classification (Fig. 4D) [34]; additionally, *SPP1^+^APOE^+^* TAM exhibited genes associated with ECM remodeling, such as *FN1*, *MMP14*, and *LGALS1* (Supplementary Fig. 3B) that are also indicative of a pro-fibrotic M2 polarization (Fig. 4D, Supplementary Fig. 3C). As the role of *FN1* in ECM signaling and EMT is paramount, we noticed that *SPP1^+^* TAM also exhibited a high EMT signature (Fig. 4E, 4F, 4G), with *FN1*, *LRP1*, *PLAUR*, and *TIMP1* highly expressed (Supplementary Fig 3B) and identified as important components leading to EMT [35,36]. Moreover, pathway analysis revealed that *SPP1^+^APOE^+^* TAM were regulated by the *HIF-1* signaling pathway (Supplementary Fig. 3C, 3D), inducing expression of *VEGFA*, *LDHA*, and *ALDOA* [37,38], which not only promote hypoxia (Fig. 4G) but is another approach to induce EMT in tumor cells [39]. We also identified *SPP1^+^APOE*^+^ TAM as major downstream components of *mTORC1* signaling (Fig. 4G, Supplementary Fig. 3C), which has important roles in macrophage polarization, tumor metabolism, and protein synthesis [40], possibly hinting that *mTORC1* is a viable candidate for inducing M2 macrophages in PDAC [41]. Survival analysis conducted in the TCGA cohort showed that the signature of these TAM led to worse survival for patients (Fig. 4H). Thus, we have established the monumental impact of these macrophages in the immuno-suppresive, pro-fibrotic niche, which we hypothesize may help support the role of *CTHRC1^+^GREM1^+^* myCAF and tumor cells further through EMT and other pro-tumor mechanisms [42]. Together, these interactions reveal the multi-faceted roles of *SPP1^+^* TAM in the PDAC TME.

### Expressions of *CTHRC1^+^GREM1^+^* CAF and *SPP1^+^APOE^+^* TAM Correspond with Worse Survival

Because of the significant contributions of *CTHCR1^+^GREM1^+^* myCAF and *SPP1^+^APOE^+^* TAM to a pro-tumorigenic TME, we investigated the potential of a synergistic relationship. To address this hypothesis, we first conducted expression plots of *CTHRC1*, *GREM1*, *SPP1*, and *APOE* in the TCGA cohort (Fig. 5A, Supplementary Fig. 4A), which were all significantly enriched in tumor samples versus control. To determine if these two groups were related, Spearman correlation (Fig. 5B) revealed a remarkable correlation between the myofibroblast and macrophage signatures (r = 0.87), while the relationship between *CTHRC1* and *SPP1* alone was also deemed significant (r = 0.25). Survival analysis of *CTHRC1* and *SPP1* gene expression in PAAD demonstrated that patients with high expression of these genes led to worse survival (Fig. 5C), demonstrating the pro-tumor functions of these genes in the clinical context. Furthermore, the combined gene signature of both groups also resulted in worse prognosis for patients, as well as being enriched in tumor populations (Fig. 5C-D). These results highlight the pro-tumorigenic and positive correlation of *CTHRC1^+^GREM1^+^*myCAF and *SPP1^+^APOE^+^* macrophages in the TME in single-cell resolution. Next, we aimed to demonstrate their relationship using spatial transcriptomics, amplifying research significance if found to be consistent with single cell transcriptomics.

**Figure 5:**
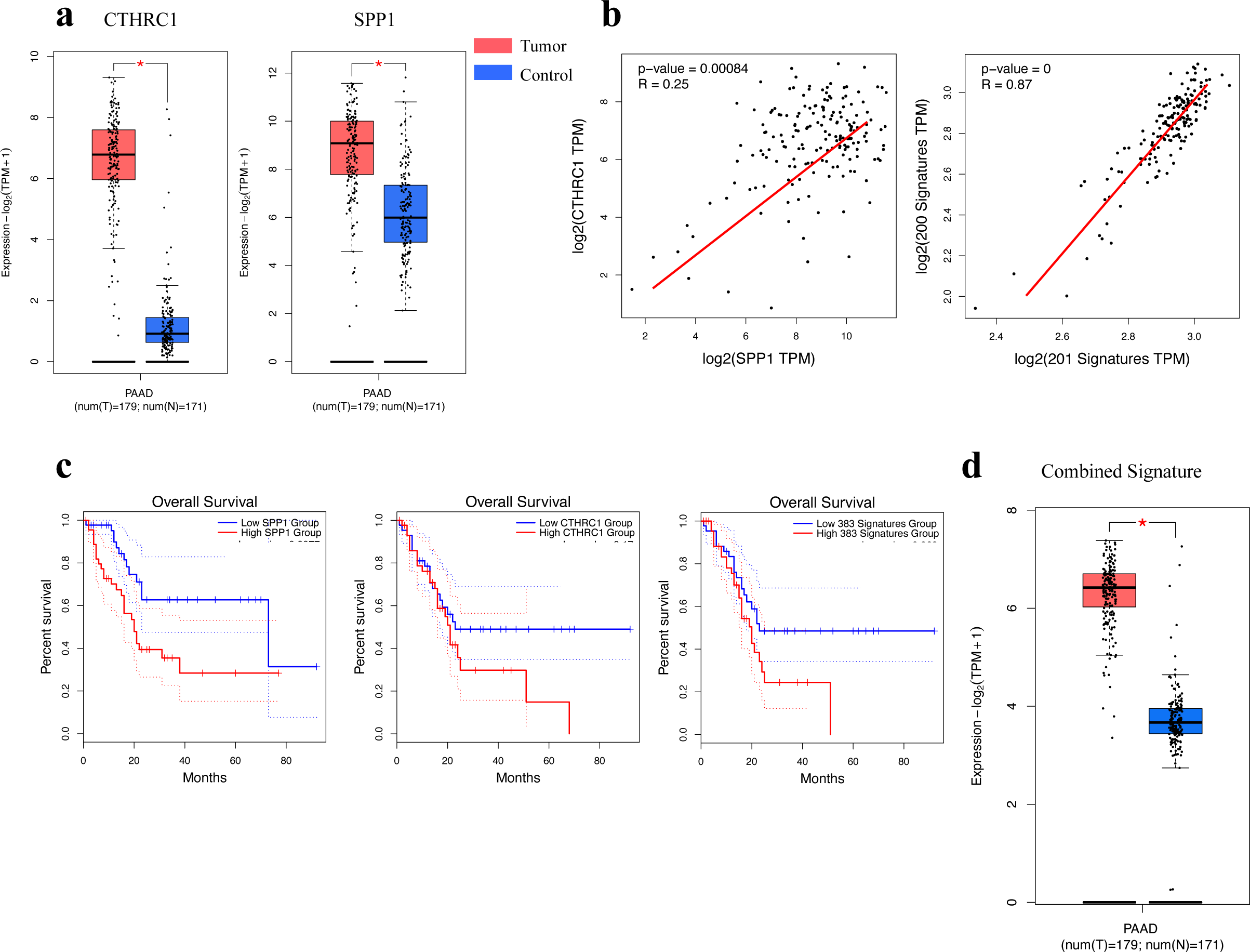
Correlation Between CTHRC1+ fibroblasts and SPP1+ macrophages lead to worse prognosis. *SPP1^+^APOE^+^* TAM and *CTHRC1^+^GREM1^+^*myCAF are positively correlated and contribute to worse prognosis in cancer patients. (A) Box plots of *CTHRC1* and *SPP1* gene expression in control versus tumor patients (TCGA cohort). (B) Spearman correlation graphs of *CTHRC1* and *SPP1*, and of the *CTHRC1^+^GREM1^+^*myCAF signature and *SPP1^+^APOE^+^* TAM signature. (C) Overall survival (OS) curves of *CTHRC1^+^GREM1^+^* myCAF, *SPP1^+^APOE^+^* TAM, and the combined signatures of both. (D) Expression boxplot of the combined signature.

### Spatial Transcriptomics Reveals Co-Localization of *CTHRC1^+^GREM1^+^* CAF and *SPP1^+^APOE^+^* TAM

To assess cell interactions in the spatial landscape, we conducted spatial transcriptomics on three public datasets acquired from PDAC patients (Fig. 6A, Supplementary Fig. 5A). A large population of ductal cells were identified based on expression of *PRSS1* and *REG1A*; populations of alpha cells (GCG) and beta cells were also observed (INS), but were all categorized under the ductal population. We also identified mixed groups, including a fibroblast/ductal population characterized by high expression of *COL1A1*, *ACTA2*, and *REG1A*; a fibroblast/malignant population expressed *GREM1*, *TIMP1*, and *TFF1*, and a mixed immune population had markers *CD3E* and *MS4A1*. In addition to normal cells of the pancreas, tumor cells were determined based on their expression of *EPCAM*, *CDH1*, and *ID1*, while an epithelial cluster was established based on the unique expression of *MUC4*, *MUC6*, and *MUC5AC*. We also noticed a substantial population of *CTHRC1^+^GREM1^+^* myofibroblasts which expressed high levels of ECM genes (*CTHRC1*, *GREM1*, *FN1*, *POSTN*), aligned with scRNAseq results (Fig. 6B, Supplementary Fig. 5B). Additionally, we found the aforementioned *SPP1^+^* macrophages, with expression of *SPP1*, *CD68*, *MARCO*, and *FN1* (Fig. 6B). We verified this by plotting the *CTHRC1^+^GREM1^+^* myCAF, *SPP1^+^APOE^+^*TAM, and EMT signatures from previous parts of our study (Fig. 6C, Supplementary Fig. 5C), which showed clear overlays, supporting the co-localization of fibroblasts and macrophages while promoting pro-tumor TME. Furthermore, the myCAF and EMT signature were almost identical, spatially confirming the presence of *CTHRC1^+^GREM1^+^* myCAF in supporting EMT. Spearman correlation between the myCAF and TAM signatures (r = 0.64) further confirmed the synergistic relationship between these two groups in the spatial landscape (Fig. 6D). Since all three groups of fibroblasts, macrophages, and tumor cells were identified in close proximity to each other (Fig. 6A), we then quantitatively visualized these correlations with a proximity enrichment heatmap (Fig. 6E), showing that *CTHRC1^+^GREM1^+^* myCAF were only in relationship to *SPP1*^+^ TAM and tumor cells, while *SPP1+* TAM were in close proximity to tumor cells and also to immune cells.

**Figure 6:**
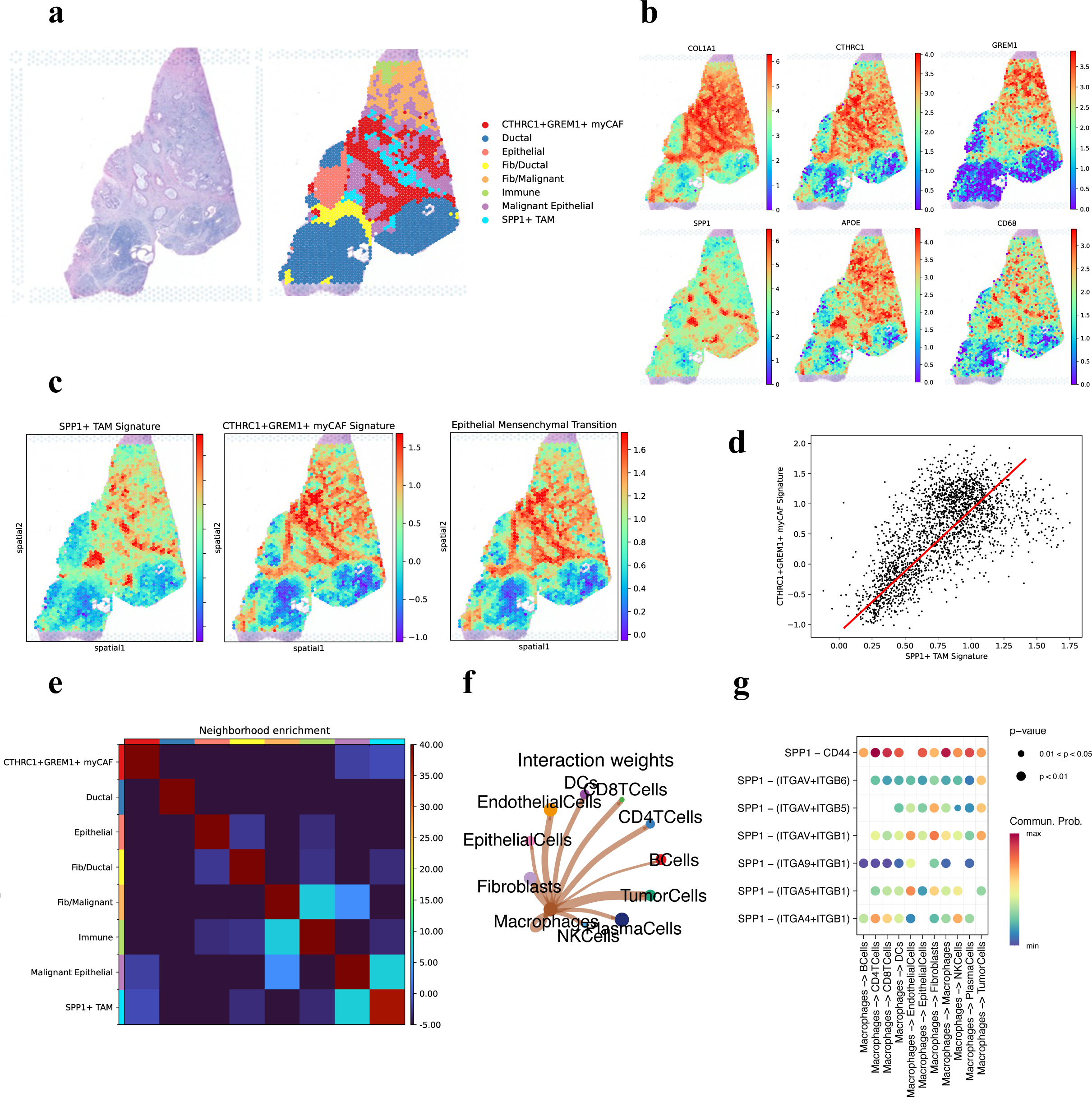
Spatial Landscape Reveals Co-Localization of fibroblasts and macrophages in TME. Spatial analysis confirms co-localization of *CTHRC1^+^GREM1^+^* myCAF and *SPP1^+^* TAM. (A) Representative histopathological H&E (FFPE) staining of pancreas tissue cross-section. UMAP of Leiden clusters shows colocalization of *CTHRC1^+^GREM1^+^* myCAF and *SPP1^+^* TAM. (B) Spatial gene expression of important marker genes. (C) Spatial enrichment of *SPP1^+^* TAM, *CTHRC1^+^GREM1^+^*myCAF, and EMT pathway signatures based on scRNAseq. (D) Spearman correlation graph of *SPP1^+^* TAM and *CTHRC1^+^GREM1^+^* myCAF signatures. (E) Neighborhood enrichment correlation matrix to quantitatively visualize spatial cellular cluster proximity. (F) Interaction weights of *SPP1*^+^ macrophage signaling to other cell types. (G) Dot plot highlights *SPP1* ligand-receptor interactions from macrophages to other cells.

### Discovering the Intricate Crosstalk Between *CTHRC1^+^GREM1^+^* myCAF, *SPP1^+^* TAM and Tumor Cells

To understand how these three populations interact, cellular ligand-receptor analysis revealed that *SPP1* expressed by macrophages interacted with tumor cells, fibroblasts, and endothelial cells the most (Fig. 6F). *SPP1* bound with integrins such as *ITGB1* and *ITGA5* (Fig. 6G), implying the role of *SPP1* in integrin signaling between fibroblasts, macrophages, and tumor cells that lead to EMT and cell-adhesion to the ECM. The *SPP1*-*CD44* axis was also significantly enriched in macrophage crosstalk (Fig. 6 G), corresponding to an important pathway for cell surface adhesion and metastasis through the activation of *PI3K/Akt* and *MAPK* signaling [42]. *SPP1^+^* macrophages also exemplified high expression of *TGFβ1* (Supplementary Fig. 5B), suggesting an increase in *TGFβ* signaling towards tumor cells and fibroblasts that directly contribute towards fibroblast recruitment, cell proliferation and EMT, although *TGFβ1* was high across all groups. Similarly, *CTHRC1^+^GREM1^+^*myCAF communicated with *SPP1^+^* macrophages and tumor cells through integrin signaling (Supplementary Fig. 5D), where collagen/integrin pairs and *FN1*/integrin pairs were prevalent. These interactions not only enhance cell proliferation and transformation [43], but could serve as a potential factor of polarization and recruitment of macrophages to the TME. Together, these interactions reveal the pathways between fibroblasts, macrophages, and tumor cells that contribute to tumor differentiation, proliferation, and worse prognosis.

## Discussion

Although there have been numerous transcriptomic studies on the TME, much work remains to be done in order to find potential drug targets, including in-depth classifications and characterization of the crosstalk between immune, stroma, and cancer cells. The inadequate therapeutic efficacy of current immunotherapeutic clinical trials underscore the importance of finding methods to overcome the immunosuppressive TME [44]. Through further analysis of single-cell RNA-seq data and using the latest approaches in spatial transcriptomics, we highlight the spatial proximity of *SPP1^+^APOE^+^* TAM and *CTHRC1^+^GREM1^+^*myCAF that concordantly contribute to the establishment of a pro-tumor TME involving immuno-regulatory, pro-fibrotic, EMT, and other mechanisms. These types of cellular compartments contain potential as cellular targets for future immune therapies.

In this study, we found that *CTHRC1^+^GREM1^+^*myCAF were enriched in tumor patients and expressed significantly higher levels of matrix-associated genes such as proteoglycans (*ACTB*, *DCN*), collagens (*COL1A2*, *COL3A1*), and matrix metalloproteinases (*MMP2*, *MMP11*, *MMP14*), indicating fibrotic properties and contributing to worse OS. To note, we did not conduct trajectory analysis and investigate the potential lineage of myCAF; however, studies done by Jin et al. and Kang et al. have revealed the function of *CTHRC1* in activating pancreatic stellate cells (PSCs) [45], which are responsible for the differentiation of PSCs into myCAF [46], suggesting a potential cycle of differentiation into the myCAF indicated in our study.

Acknowledging the *α-SMA^+/-^* classification of myCAF activation, our analysis revealed that *α-SMA* was not a defining gene of our myCAF population, and we suggest that *CTHRC1* may serve as a potential future marker for myCAF in PDAC and other solid tumor cancers due to its increased expression and positive contribution to the carcinogenesis of other cancers, such as stomach, liver, colon, and breast [47,48]. However, future studies will need to be conducted not only in PDAC, but in other cancers to validate *CTHRC1* as a defining marker in myCAF.

Recent research has been conducted to investigate whether targeting myCAF can lead to better outcomes. Nevertheless, reducing myCAF by targeting *α-SMA* in the tumor led to reduced survival rates in both humans and mice, due to the duality of extensive ECM protein deposition as an immune-physical barrier, restricting immune cell accessibility to cancer cells, but simultaneously restricting tumor growth to an extent [8]. However, our results highlight other *α-SMA^+^* populations such as pericytes and smooth muscle cells that potentially could have been inadvertently targeted in their study. These results emphasize the need to thoroughly characterize and target specific fibroblast populations in the TME as therapeutic targets. Here, the next focus of our study looks into the roles of *SPP1* and *SPP1^+^* macrophages as alternate targets. We not only identified two major groups of *SPP1^+^*macrophages that are anti-inflammatory M2 and pro-fibrotic like, but they also contribute to hypoxia and EMT, aiding *CTHRC1^+^GREM1^+^*myCAF in a synergistic relationship that leads to worse prognosis. Recent studies have indicated *SPP1* in cancers as a marker of pro-tumor macrophages and cancer cells, leading to worse prognosis for patients [42]. Moreover, Eun et al. [49] demonstrated the role of *SPP1* as a prognostic marker in hepatocellular carcinoma fibroblasts, leading to direct chemoresistance against tyrosine kinase inhibitors while driving EMT. Most importantly, the multifunctional presence of *SPP1* not only promotes activation on myCAF [50], but also drives EMT and chemoresistance to anti-*PDL1* therapy through the *STAT3* and *PI3K/AKt-mTOR* pathways [49,51,52].

In-vivo studies have highlighted how *SPP1^-^* knockout mice have been shown to lead to decreased M0/M2 infiltration, a decrease in fibrosis, and exhibit positive results in OS [53,54]; despite these encouraging results, no current human clinical studies inhibiting *SPP1* have been conducted on cancer. We found that the pro-fibrotic *SPP1^+^* matrisome-associated macrophages (MAM) described by Ouyang et al. [55] corresponds with the signature of *SPP1^+^APOE^+^*macrophages in this study, confirming the role of *SPP1^+^* macrophages in ECM development. Moreover, Fabre et al. [15] characterized a broad population of *SPP1^+^CD9^+^TREM2^+^*macrophages in liver and lung fibrotic diseases, while demonstrating the roles of type-3 inflammation cytokines *GM-CSF* and *IL-17A* in inducing these macrophages in-vitro and in-vivo. Importantly, this population matched our scRNAseq *SPP1^+^* TAM signature (*SPP1*, *GPNMB*, *CD63*, *FABP5*) precisely, suggesting that the presence and functions of their TAM can also be applied to PDAC. Our *SPP1^+^* TAM population also expressed *CD68* but had low expression of *CD206* and *CD163*, which differed slightly from another study of *SPP1^+^*macrophages in colorectal cancer that had high *CD206* expression [56]. Altogether, these studies highlight the role of *SPP1^+^* TAM in pro-tumor fibrosis; we further describe the functions of these cells in regard to their relationship with fibroblasts and tumor cells.

To date, few pieces of literature have investigated the relationship between *SPP1^+^* macrophages and myCAF; however, the emphasis on a positive correlation is unmistakably clear. Qi et al. [56] first did an in-depth study on the network between *FAP^+^*CAF and *SPP1^+^* TAM in colorectal cancer, proving the synergistic and spatial correlations of these groups, although their characterization of *SPP1^+^* TAM in the study suggested a M1 pro-inflammatory phenotype regulated by *STAT1*. Hoeft et al. [57] determined *CXCL4* as a driving factor behind the activation of *SPP1^+^* TAM, concurrently leading to activation of myCAF. Our results extend these implications to PDAC, highlighting the importance of the spatially correlated crosstalk between *SPP1^+^APOE^+^* TAM and *CTHRC1^+^GREM1^+^*myCAF that drive fibrosis, immunosuppression, and EMT in PDAC and potentially other cancers. However, more work is needed to describe in particular how these groups interact with T-cells and other anti-tumor responses. Considering the significance of these results in furthering our understanding of macrophage-fibroblast communication, these populations serve as potential therapeutic targets for not only PDAC, but applicable in a wide range of cancers and fibrotic diseases.

## Methods

### Data Collection

Altogether, five single-cell RNA-seq datasets were downloaded from the Gene Expression Omnibus (GSE242230, GSE154778, GSE155698, GSE205354, GSE212966), and one spatial dataset (GSE211895) were acquired for analysis. 63 patients were in this study, including adjacent normal (n=6), primary PDAC (n=51), and metastasis (n=6), who were in various stages of cancer progression (mostly stage 3 or 4, or taken after patient mortality).

### Preprocessing, Dimension Reduction and Clustering

All analysis was conducted in the jupyter notebook software using *scanpy* python package. Low quality cells and/or genes were filtered by gene counts and gene expression with the following restrictions: (1) cells less than 500 total gene counts, (2) genes found in less than three cells, (3) cells with over 100,000 total counts. Doublet removal was not performed during this step and subsequently removed manually when annotating clusters. Cell counts were normalized to 10,000 and logarithmized. Principal components (PC) were calculated based on variable genes using the *sc.pp.highly_variable_genes* function. We then used *Harmony* for batch correction before finding the closest 30 neighbors using *sc.pp.neighbors*. Cells were projected onto a Uniform Manifold Approximation Projection (UMAP) and clustered using the Leiden algorithm [58]. This was the first of two clusterings that we did to identify major cell types; the second enabled us to have a comprehensive overview of specific individual subtypes in each cell type. Major cell clusters were annotated based on the top 200 differential gene expression and manually confirmed (Supplementary Table 1).

### Proportion Comparison and Statistical Methods

To compare proportions of tumor versus adjacent normal in each cell subtype, we calculated the percentage using (number of cells in subtype)/(total number of cells in patient) *100 separately for both conditions. A *matplotlib* bar plot was used to visualize data, and the Mann-Whitney U-Test and Welch’s T-Test from the *statannotations* package were used to compare means (a p-value of less than 0.05 considered statistically significant).

### Differential Gene Analysis

The top differential genes of individual subtypes were calculated using *sc.tl.rank_genes_groups*, and p-values were obtained using the non-parametric Wilcoxon rank-sum test (Supplementary Table 1). For visualization, we plotted heatmaps and volcano plots (https://github.com/mousepixels/sanbomics) to show DE genes in comparison to the rest of the data. The top 200 DE genes were used to label clusters and GSEA.

### Gene Signature Rankings and GSEA

Signature gene lists were obtained from GSEA (https://www.gsea-msigdb.org/gsea/msigdb/human/genesets.jsp) and through literature (Supplementary Table 2). Gene signature scores were calculated using *sc.tl.score_genes* function and visualized on the UMAP embedding. We ran GSEA using GSEApy on all significant cell subtypes with MSigDB_Hallmark and KEGG to generate figures, with a p-value and false discovery rate of less than 0.05 considered significant. In addition, we utilized the Enrichr (https://maayanlab.cloud/Enrichr/) analysis method to acquire different rankings of unregulated pathways and mechanisms of each gene list.

### Survival Analysis

Survival analysis and Kaplan-Meier survival curves of overall survival and disease free survival (RFS) was conducted in GEPIA (http://gepia2.cancer-pku.cn/#survival) using all PAAD cohorts with a 95% confidence interval. Hazard ratio was calculated using the Cox proportional hazards model. Both individual genes and gene signatures of differentially expressed genes were used as input.

### Spatial Transcriptomics Analysis

Three spatial immunofluorescent cross-tissue slides were acquired from public datasets (GSE211895) using the Visium platform from 10X Genomics. Principal components were calculated based on the top variable genes per spot using *scanpy*. Dimension reduction, neighbors, and leiden clustering were performed to generate different clusters of spots. The top 200 DE genes were exported and utilized to perform manual annotations. UMAP of individual gene features were conducted in the same manner as single cell data, as well as cell signatures taken from scRNAseq data. *Squidpy* package was used to project a neighborhood enrichment heatmap of all clusters to confirm relationships between clusters. Spearman correlation was calculated using *scipy.* We also utilized Ye Labs’ spatial analysis to cross-reference our results (http://labwebsite.yelab.site:1234/<u>#!/</u>).

### Ligand-Receptor Communications

To extrapolate cell-cell communication between cell subtypes of interest (*SPP1*^+^ TAM, *CTHRC1^+^GREM1^+^* myCAF, and tumor cells), we used *LIANA* (https://github.com/saezlab/liana-py) as a framework to conduct ligand-receptor analysis on single-cell and spatial data using CellChat and CellPhoneDB. The *rank_aggregate* method averaged specificity and magnitude across all methods. We also used the Ye Labs spatial analysis to pull out the top specific interactions to each group for each gene.

## Acknowledgements

The authors would like to thank Brian Kondek and Hanna Shebert for their contributions and feedback during the revision process, and a special thanks to Dr. Li Li for research guidance.

## Author Contributions

E.L. was the main contributor and researcher of the project, including analysis, data interpretation, and results. H.C.C. assisted with final manuscript preparation and revision. S.M. assisted with results and final manuscript preparation, including introduction and discussion.

## Data Availability

Public scRNA-seq datasets conducted in this study are accessible through the Gene Expression Omnibus, including GSE242230 (https://www.ncbi.nlm.nih.gov/geo/query/acc.cgi?acc=GSE242230), GSE154778 (https://www.ncbi.nlm.nih.gov/geo/query/acc.cgi?acc=GSE154778), GSE155698 (https://www.ncbi.nlm.nih.gov/geo/query/acc.cgi?acc=GSE155698), GSE205354 (https://www.ncbi.nlm.nih.gov/geo/query/acc.cgi?acc=GSE205354), and GSE212966 (https://www.ncbi.nlm.nih.gov/geo/query/acc.cgi?acc=GSE212966). The TCGA data (https://gdac.broadinstitute.org/#) was analyzed through GEPIA (http://gepia2.cancer-pku.cn/#index). Spatial transcriptomics data are available in GSE211895 (https://www.ncbi.nlm.nih.gov/geo/query/acc.cgi?acc=GSE211895), and analyzed with Ye Labs TME website (http://labwebsite.yelab.site:1234/#!/) in conjunction with our own methods.

Code availability is accessible upon request.

## Ethics

The authors declare no competing interests.

