## Supplementary Figures for "*CTHRC1*^+^ Fibroblasts and *SPP1*^+^ Macrophages Synergistically Contribute to Pro-Tumorigenic Tumor Microenvironment in Pancreatic Ductal Adenocarcinoma"

### Fig. 1 Supplementary

## a

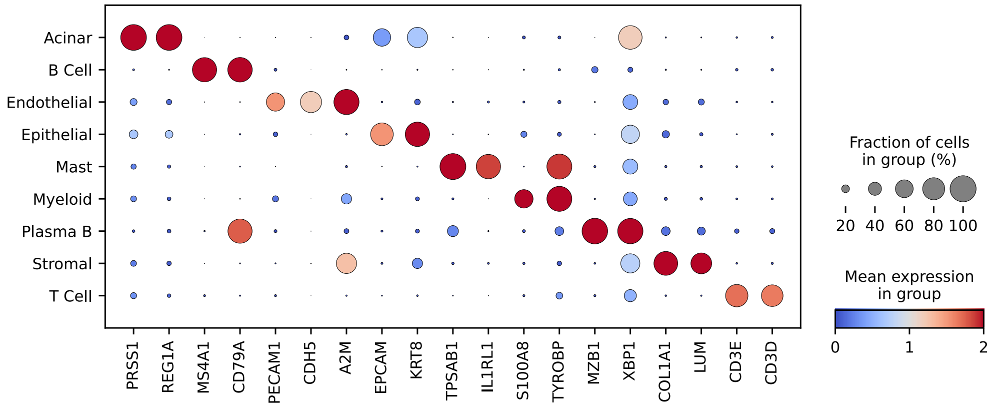


(A) Dot plot of canonical marker genes in major cell clusters.

### Fig. 2 Supplementary

## a

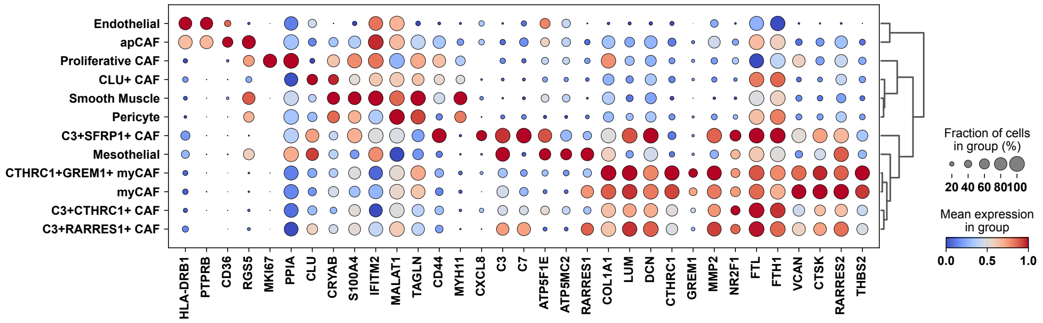


## b

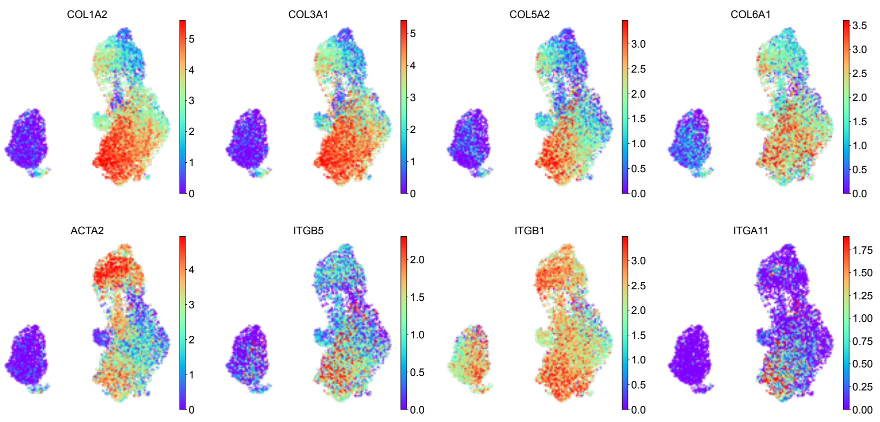

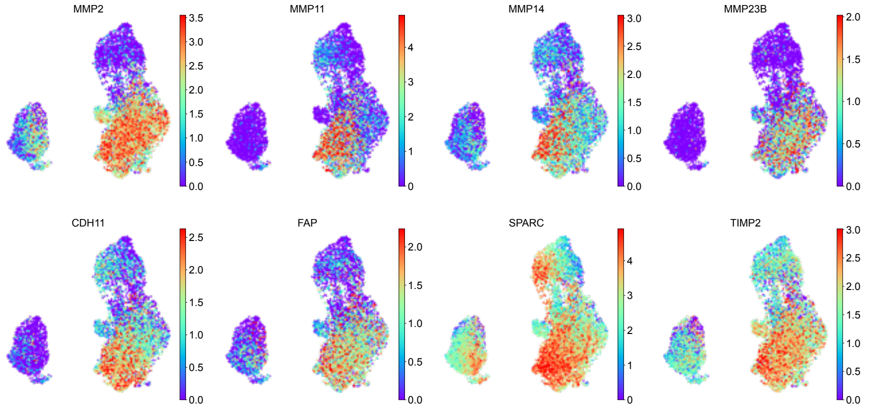
MMP2
MMP11
MMP14
MMP23B
CDH11
FAP
SPARC
TIMP2

c

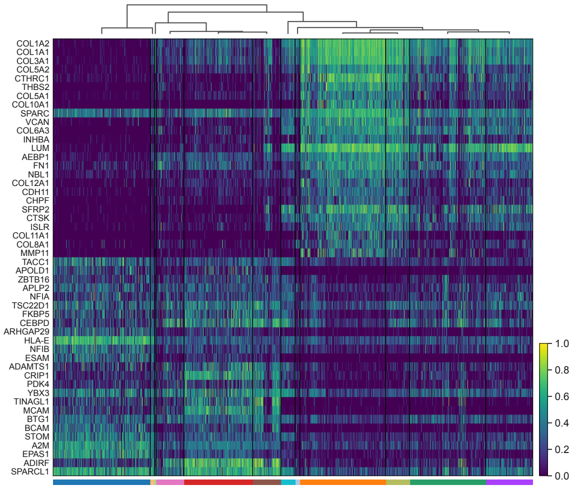
COL1A2
COL1A1
COL3A1
COL5A2
CTHRC1
THBS2
COL5A1
COL10A1
SPARC
VCAN
COL6A3
INHBA
LUM
AEBP1
FN1
NBL1
COL12A1
CDH11
CHPF
SFRP2
CTSK
ISLR
COL11A1
COL8A1
MMP11
TACC1
APOLD1
ZBTB16
APLP2
NFIA
TSC22D1
FKBP5
CEBPD
ARHGAP29
HLA-E
NFIB
ESAM
ADAMTS1
CRIP1
PDK4
YBX3
TINAGL1
MCAM
BTG1
BCAM
STOM
A2M
EPAS1
ADIRF
SPARCL1

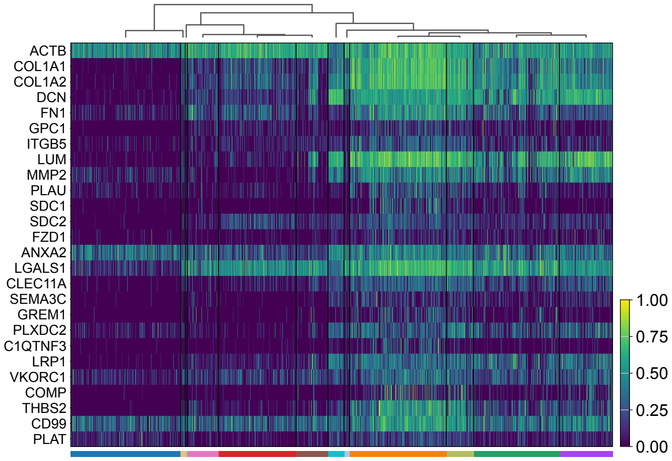
ACTB
COL1A1
COL1A2
DCN
FN1
GPC1
ITGB5
LUM
MMP2
PLAU
SDC1
SDC2
FZD1
ANXA2
LGALS1
CLEC11A
SEMA3C
GREM1
PLXDC2
C1QTNF3
LRP1
VKORC1
COMP
THBS2
CD99
PLAT
1.00
0.75
0.50
0.25
0.00

d

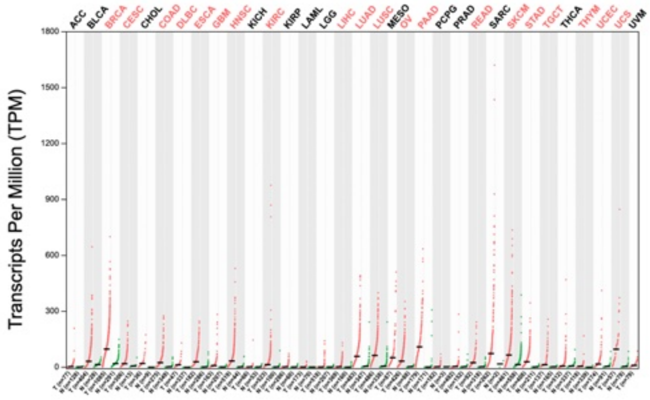
Transcripts Per Million (TPM)

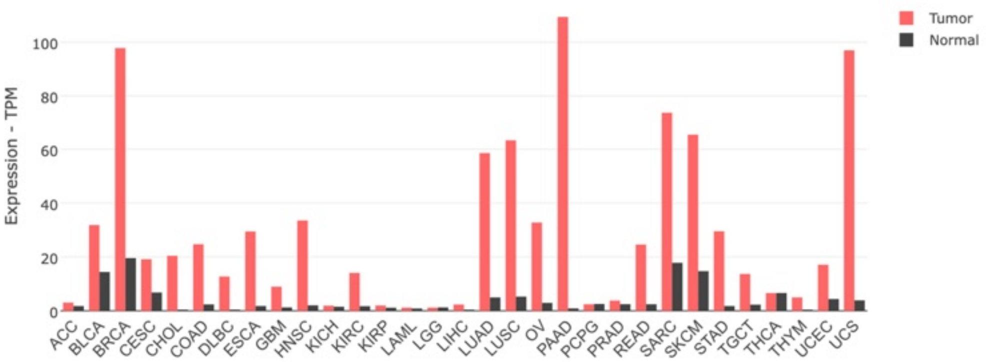
Tumor
Normal
Expression - TPM
ACC
BLCA
BRCA
CESC
CHOL
COAD
DLBC
ESCA
GBM
HNSC
KICH
KIRC
KIRP
LAML
LGG
LIHC
LUAD
LUSC
OV
PAAD
PCPG
PRAD
READ
SARC
SKCM
STAD
TGCT
THCA
THYM
UCEC
UCS

e

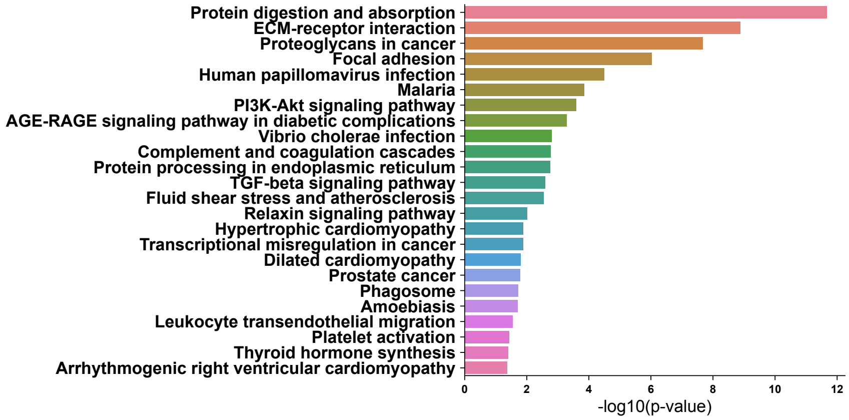


f

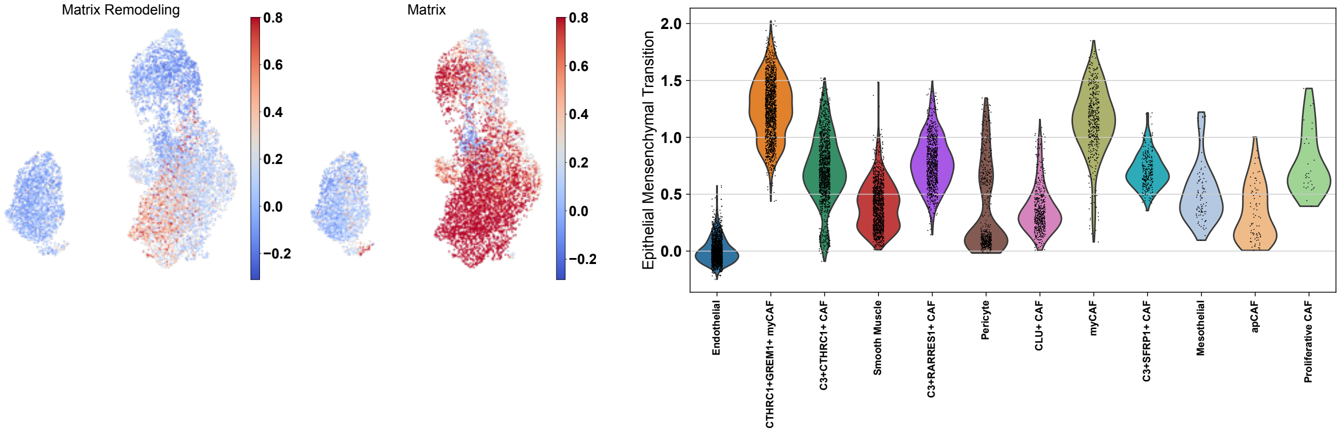


g

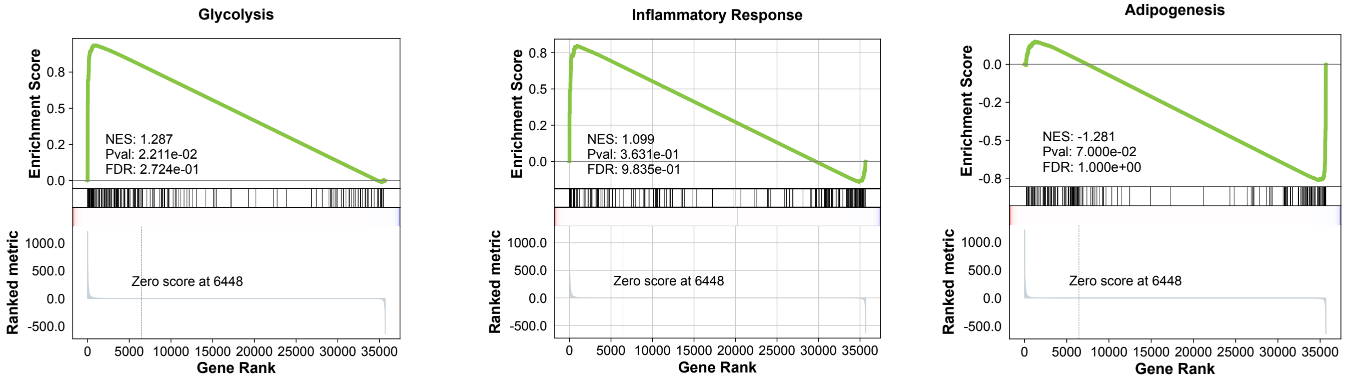


(A) Dot plot of stromal cell populations in the TME. (B) UMAP of matrix-associated genes enriched in *CTHRC1*$^{+}$*GREM1*$^{+}$ myCAF. (C) Heatmaps of top 25 DE genes enriched in *CTHRC1*$^{+}$*GREM1*$^{+}$ myCAF and proteoglycans. (D) TPM Expression of *CTHRC1* in the TCGA cohort, compared with all cancers. TPM of *CTHRC1* in tumor versus control in all cancers. (E) UMAP of matrix, matrix remodeling, and TGBβ signaling enrichment signatures. (F) Bar plot of upregulated KEGG enrichment pathways in *CTHRC1*$^{+}$*GREM1*$^{+}$ myCAF using Enrichr. (G) Additional GSEA plots of upregulated pathways in *CTHRC1*$^{+}$*GREM1*$^{+}$ myCAF.

### Fig. 3 Supplementary

## a

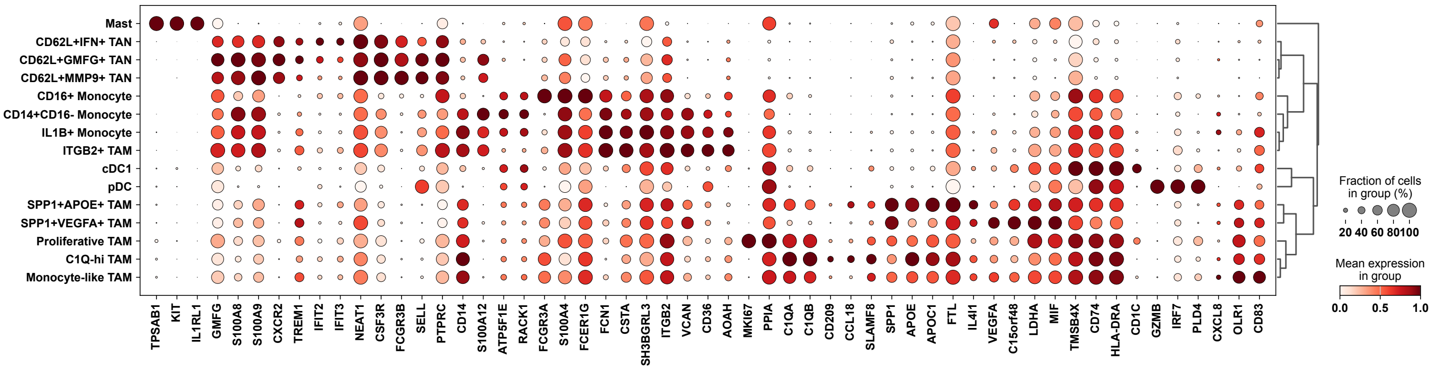


## b

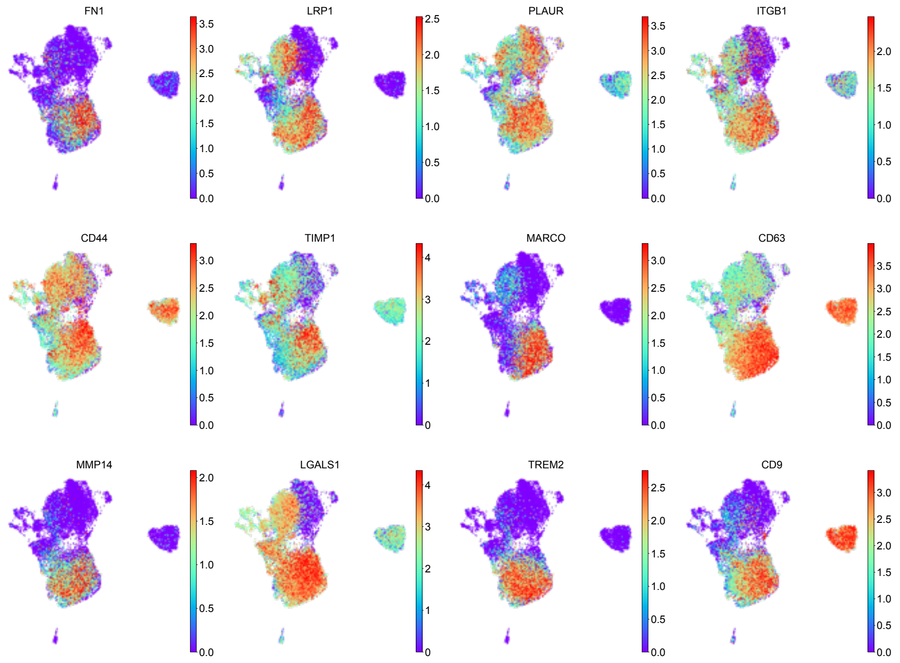


## c

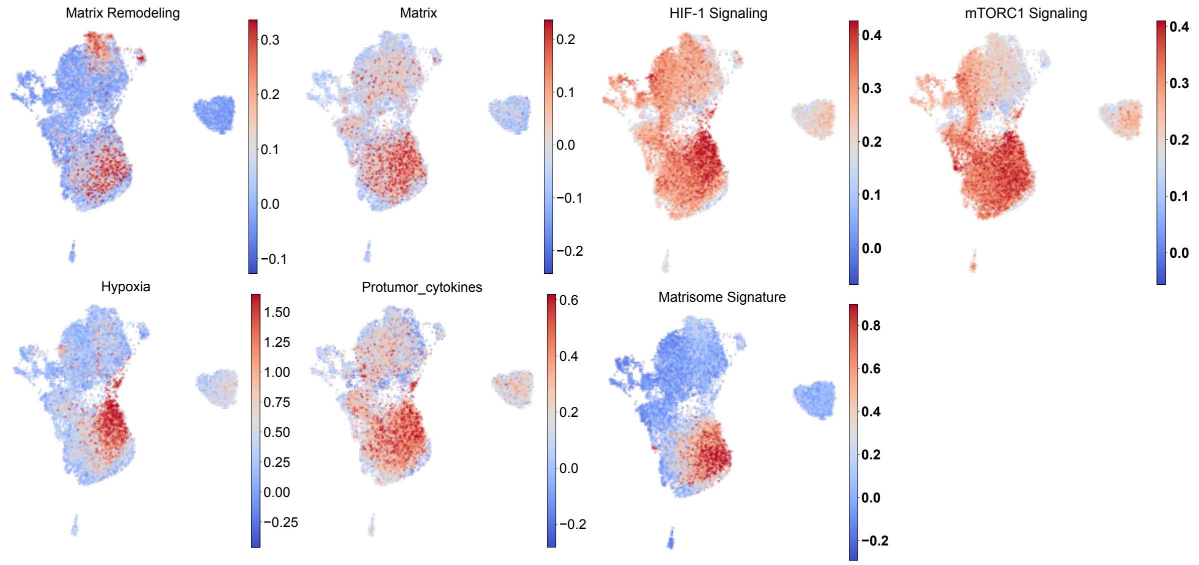

d

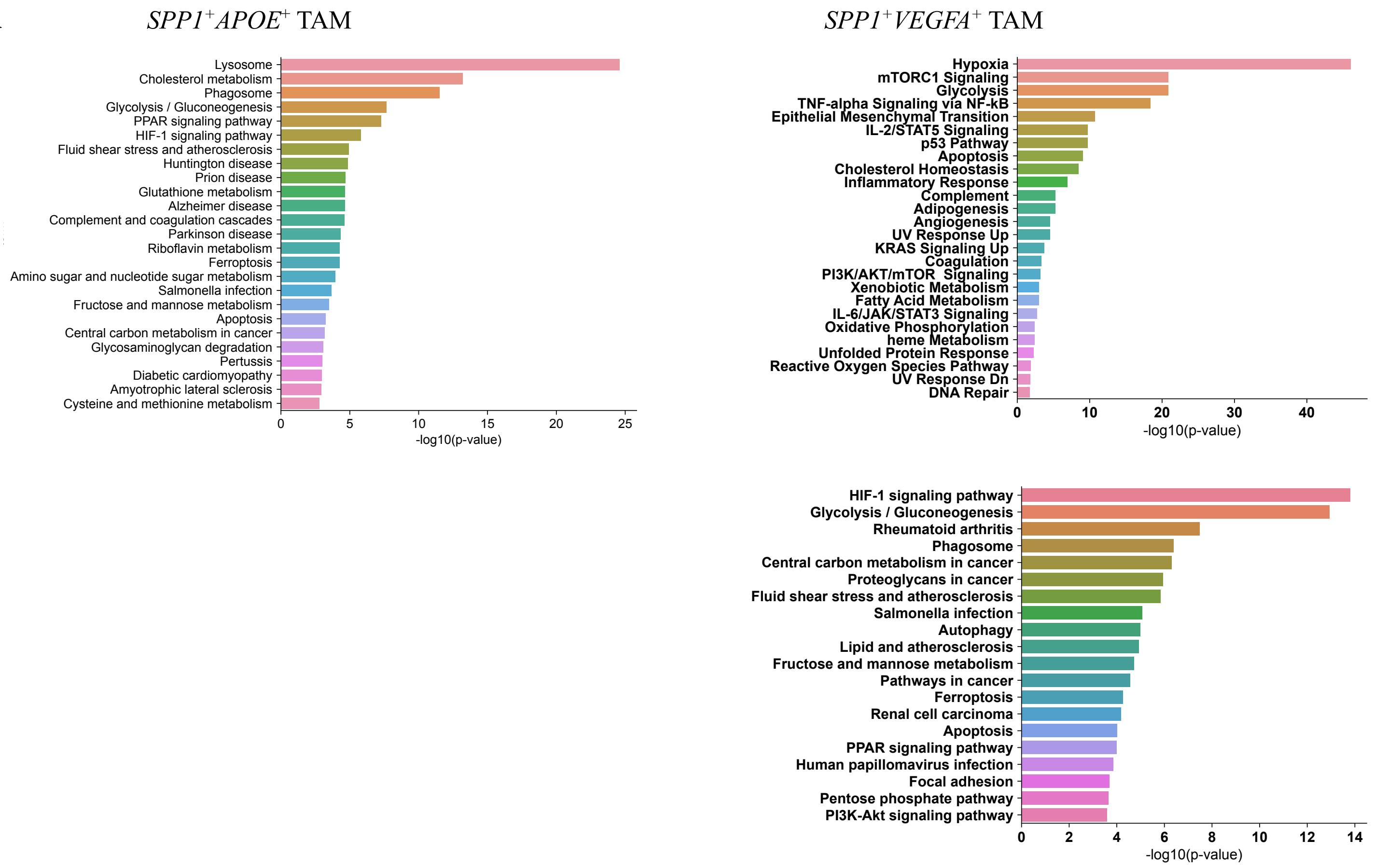


e

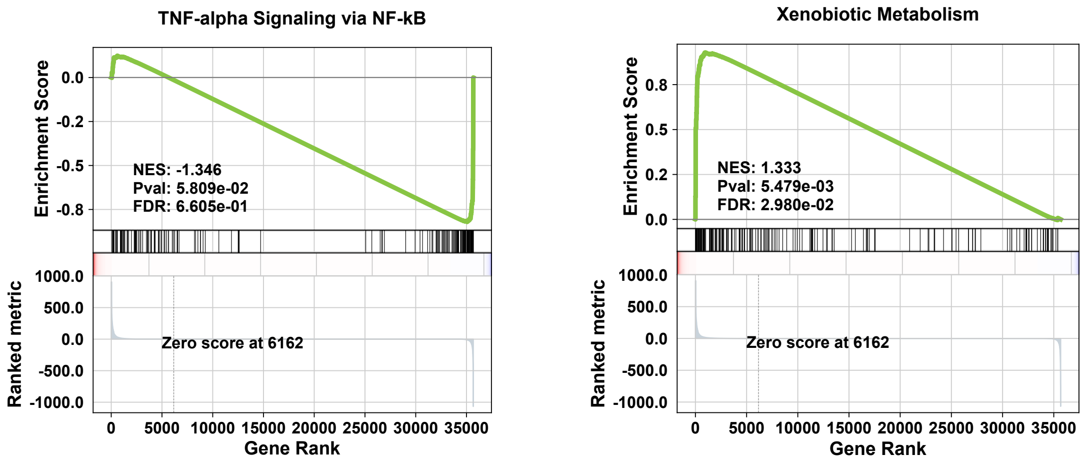


(A) Dot plot of myeloid cell populations in the TME. (B) UMAP of enriched genes expressed by *SPP1*$^{+}$ macrophages. (C) UMAP of matrix, matrix remodeling, hypoxia, M1 and M2 signatures, and pro-tumor cytokine enrichment signatures. (D) Bar plot of upregulated KEGG pathways in *SPP1*$^{+}$*APOE*$^{+}$ TAM. Bar plots of upregulated MSigDB and KEGG pathways in *SPP1*$^{+}$*VEGFA*$^{+}$ TAM. (E) GSEA plots of enriched pathways in *SPP1*$^{+}$*APOE*$^{+}$ TAM.

### Fig 4 Supplementary

**a**

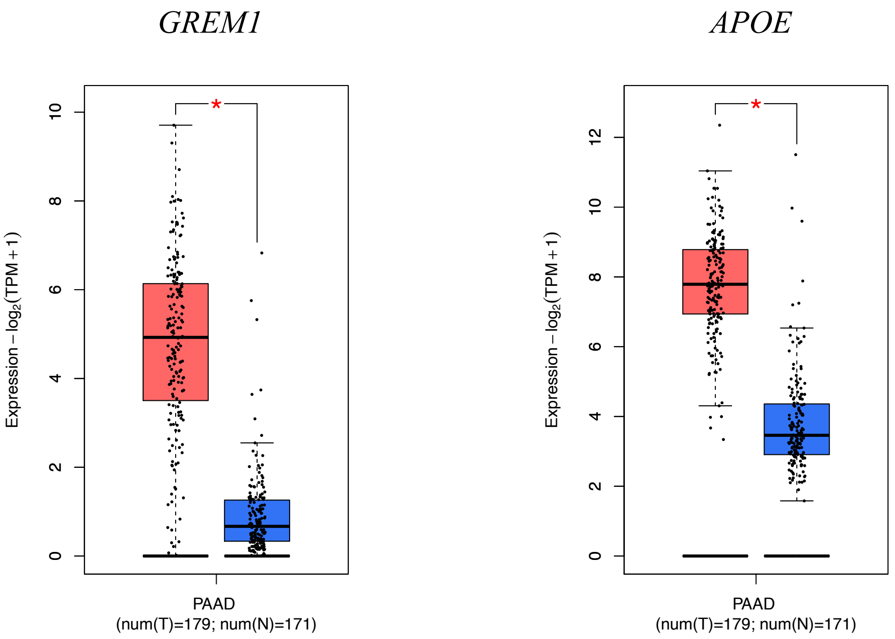


(A) Expression box plots of *GREM1* and *APOE* gene expression in control versus tumor patients (TCGA cohort).

### Fig. 5 Supplementary

**a**

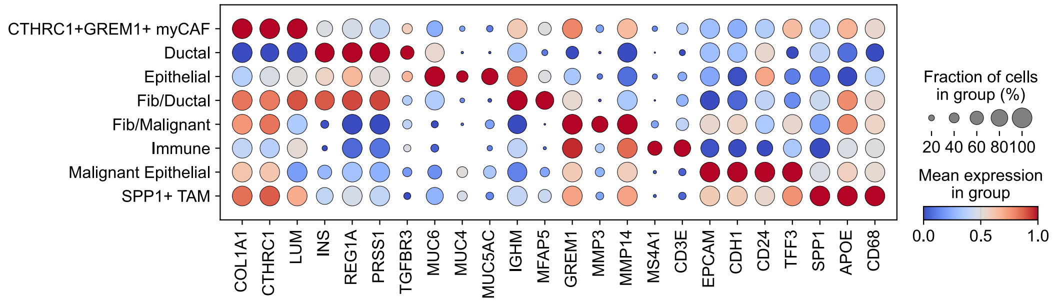


**b**

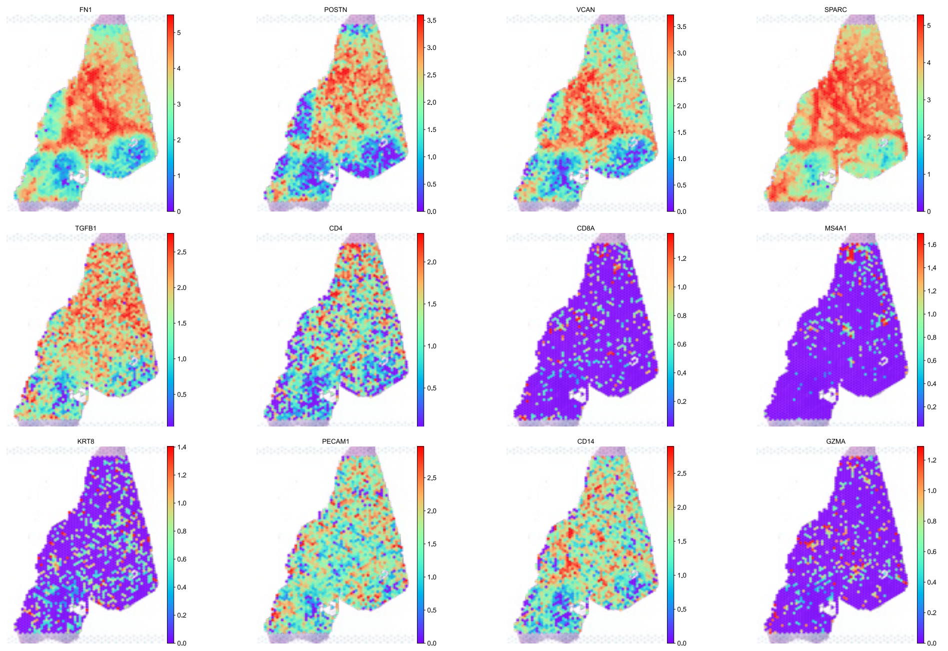

**c**

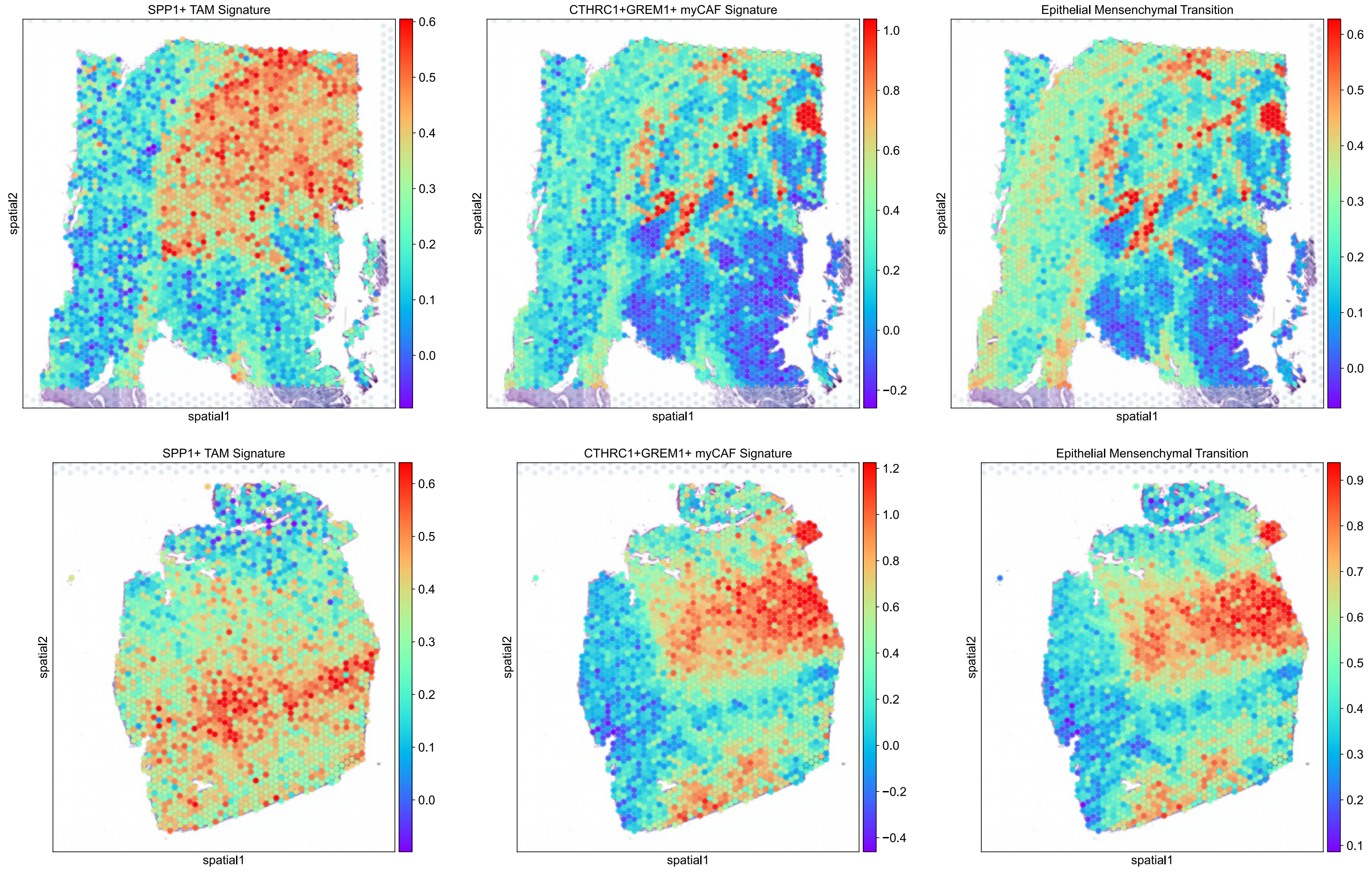
SPP1+ TAM Signature
spatial2
spatial1
0.6
0.5
0.4
0.3
0.2
0.1
0.0
CTHRC1+GREM1+ myCAF Signature
spatial2
spatial1
1.0
0.8
0.6
0.4
0.2
0.0
−0.2
Epithelial Mensenchymal Transition
spatial2
spatial1
0.6
0.5
0.4
0.3
0.2
0.1
0.0
SPP1+ TAM Signature
spatial2
spatial1
0.6
0.5
0.4
0.3
0.2
0.1
0.0
CTHRC1+GREM1+ myCAF Signature
spatial2
spatial1
1.2
1.0
0.8
0.6
0.4
0.2
0.0
−0.2
−0.4
Epithelial Mensenchymal Transition
spatial2
spatial1
0.9
0.8
0.7
0.6
0.5
0.4
0.3
0.2
0.1

**d**

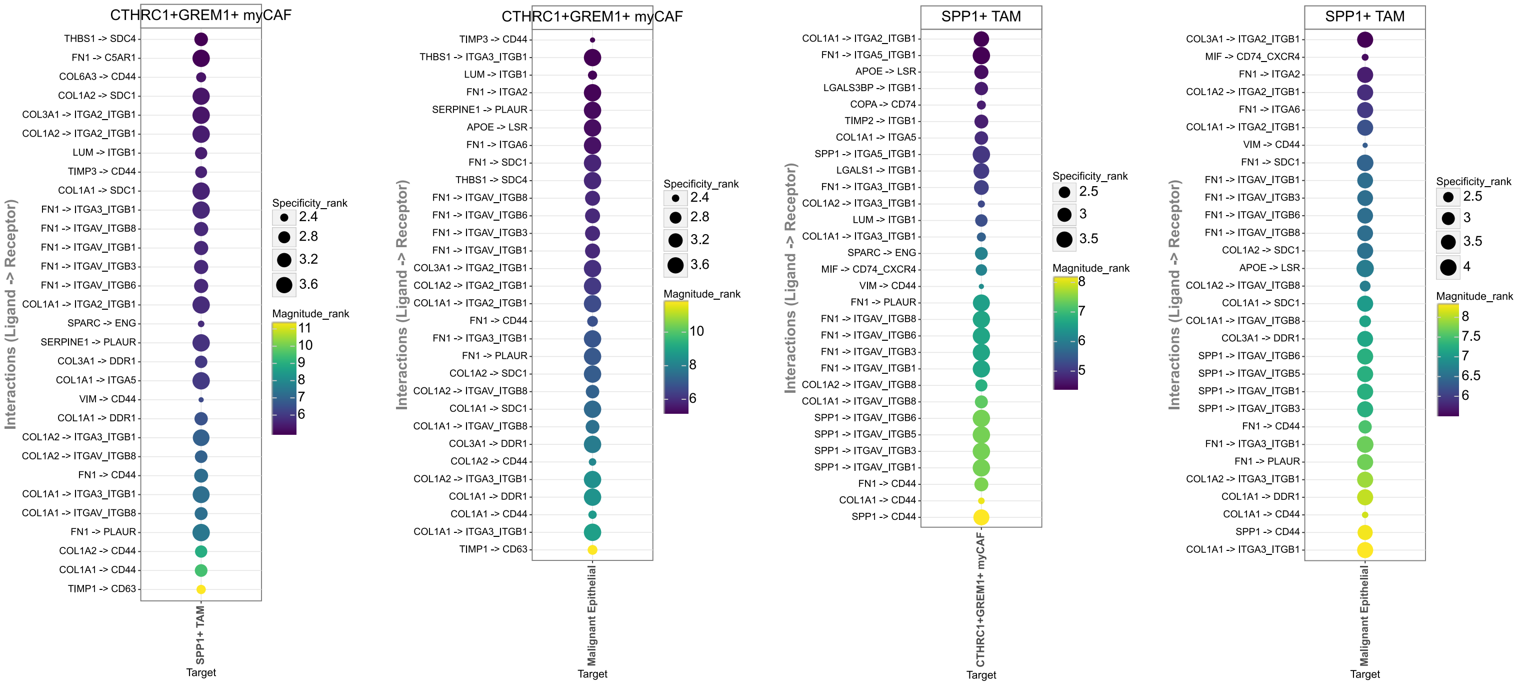
CTHRC1+GREM1+ myCAF
Interactions (Ligand -> Receptor)
THBS1 -> SDC4
FN1 -> C5AR1
COL6A3 -> CD44
COL1A2 -> SDC1
COL3A1 -> ITGA2_ITGB1
COL1A2 -> ITGA2_ITGB1
LUM -> ITGB1
TIMP3 -> CD44
COL1A1 -> SDC1
FN1 -> ITGA3_ITGB1
FN1 -> ITGAV_ITGB8
FN1 -> ITGAV_ITGB1
FN1 -> ITGAV_ITGB3
FN1 -> ITGAV_ITGB6
COL1A1 -> ITGA2_ITGB1
SPARC -> ENG
SERPINE1 -> PLAUR
COL3A1 -> DDR1
COL1A1 -> ITGA5
VIM -> CD44
COL1A1 -> DDR1
COL1A2 -> ITGA3_ITGB1
COL1A2 -> ITGAV_ITGB8
FN1 -> CD44
COL1A1 -> ITGA3_ITGB1
COL1A1 -> ITGAV_ITGB8
FN1 -> PLAUR
COL1A2 -> CD44
COL1A1 -> CD44
TIMP1 -> CD63
SPP1+ TAM
Target
Specificity_rank
2.4
2.8
3.2
3.6
Magnitude_rank
11
10
9
8
7
6
CTHRC1+GREM1+ myCAF
Interactions (Ligand -> Receptor)
TIMP3 -> CD44
THBS1 -> ITGA3_ITGB1
LUM -> ITGB1
FN1 -> ITGA2
SERPINE1 -> PLAUR
APOE -> LSR
FN1 -> ITGA6
FN1 -> SDC1
THBS1 -> SDC4
FN1 -> ITGAV_ITGB8
FN1 -> ITGAV_ITGB6
FN1 -> ITGAV_ITGB3
FN1 -> ITGAV_ITGB1
COL3A1 -> ITGA2_ITGB1
COL1A2 -> ITGA2_ITGB1
COL1A1 -> ITGA2_ITGB1
FN1 -> CD44
FN1 -> ITGA3_ITGB1
FN1 -> PLAUR
COL1A2 -> SDC1
COL1A2 -> ITGAV_ITGB8
COL1A1 -> SDC1
COL1A1 -> ITGAV_ITGB8
COL3A1 -> DDR1
COL1A2 -> CD44
COL1A2 -> ITGA3_ITGB1
COL1A1 -> DDR1
COL1A1 -> CD44
COL1A1 -> ITGA3_ITGB1
TIMP1 -> CD63
Malignant Epithelial
Target
Specificity_rank
2.4
2.8
3.2
3.6
Magnitude_rank
10
8
6
SPP1+ TAM
Interactions (Ligand -> Receptor)
COL1A1 -> ITGA2_ITGB1
FN1 -> ITGA5_ITGB1
APOE -> LSR
LGALS3BP -> ITGB1
COPA -> CD74
TIMP2 -> ITGB1
COL1A1 -> ITGA5
SPP1 -> ITGA5_ITGB1
LGALS1 -> ITGB1
FN1 -> ITGA3_ITGB1
COL1A2 -> ITGA3_ITGB1
LUM -> ITGB1
COL1A1 -> ITGA3_ITGB1
SPARC -> ENG
MIF -> CD74_CXCR4
VIM -> CD44
FN1 -> PLAUR
FN1 -> ITGAV_ITGB8
FN1 -> ITGAV_ITGB6
FN1 -> ITGAV_ITGB3
FN1 -> ITGAV_ITGB1
COL1A2 -> ITGAV_ITGB8
COL1A1 -> ITGAV_ITGB8
SPP1 -> ITGAV_ITGB6
SPP1 -> ITGAV_ITGB5
SPP1 -> ITGAV_ITGB3
SPP1 -> ITGAV_ITGB1
FN1 -> CD44
COL1A1 -> CD44
SPP1 -> CD44
CTHRC1+GREM1+ myCAF
Target
Specificity_rank
2.5
3
3.5
Magnitude_rank
8
7
6
5
SPP1+ TAM
Interactions (Ligand -> Receptor)
COL3A1 -> ITGA2_ITGB1
MIF -> CD74_CXCR4
FN1 -> ITGA2
COL1A2 -> ITGA2_ITGB1
FN1 -> ITGA6
COL1A1 -> ITGA2_ITGB1
VIM -> CD44
FN1 -> SDC1
FN1 -> ITGAV_ITGB1
FN1 -> ITGAV_ITGB3
FN1 -> ITGAV_ITGB6
FN1 -> ITGAV_ITGB8
COL1A2 -> SDC1
APOE -> LSR
COL1A2 -> ITGAV_ITGB8
COL1A1 -> SDC1
COL1A1 -> ITGAV_ITGB8
COL3A1 -> DDR1
SPP1 -> ITGAV_ITGB6
SPP1 -> ITGAV_ITGB5
SPP1 -> ITGAV_ITGB1
SPP1 -> ITGAV_ITGB3
FN1 -> CD44
FN1 -> ITGA3_ITGB1
FN1 -> PLAUR
COL1A2 -> ITGA3_ITGB1
COL1A1 -> DDR1
COL1A1 -> CD44
SPP1 -> CD44
COL1A1 -> ITGA3_ITGB1
Malignant Epithelial
Target
Specificity_rank
2.5
3
3.5
4
Magnitude_rank
8
7.5
7
6.5
6

(A) Dot plot of spatial cell populations in the TME (Patient 1). (B) Spatial expression plots of matrix-associated genes and immune cell canonical markers. (C) Spatial enrichment of *SPP1*$^{+}$*APOE*$^{+}$ TAM, *CTHRC1*$^{+}$*GREM1*$^{+}$ myCAF, and EMT signatures in patients 2 and 3. (D) Top Liana ligand-receptor interactions of *CTHRC1*$^{+}$*GREM1*$^{+}$ myCAF and *SPP1*$^{+}$ TAM.
